# In V4, pair-wise synchrony and correlations depend on the structure of the population code

**DOI:** 10.1101/645135

**Authors:** Veronika Koren, Ariana R. Andrei, Ming Hu, Valentin Dragoi, Klaus Obermayer

## Abstract

In visual areas of primates, neurons activate in parallel while the animal is engaged in a behavioral task. In this study, we examine the structure of the population code while the animal performs delayed match to sample task on complex natural images. The macaque monkeys visualized two consecutive stimuli that were either the same or different, while recorded with laminar arrays across the cortical depth in cortical areas V1 and V4. We decoded correct choice behavior from neural populations of simultaneously recorded units. Utilizing decoding weights, we divide neurons in most informative and less informative, and show that most informative neurons in V4, but not in V1, are more strongly synchronized, coupled and correlated than less informative neurons. As neurons are divided in two coding pools according to their coding preference, in V4, but not in V1, spiking synchrony, coupling and correlations within the coding pool are stronger than across coding pools.

**Highlights:**

- In a match-to-sample visual task, responses of neural populations in V1 and in V4 predict the stimulus class better than chance.
- In V4, informative neurons are more strongly coupled, correlated and synchronized than less informative neurons.
- In V4, neurons are more strongly coupled, correlated and synchronized within coding pools compared to across coding pools.
- Correlations within coding pools harm the performance of the classifier in both V1 and V4.

## Introduction

The general understanding is that sensory neurons respond to features of sensory stimuli and transmit this information downstream. Downstream neurons read-out activity of projecting neurons and use it for generating a signal that is highly correlated with the behavioral choice of the animal ([Shadlen and Newsome, 2001]). Traditionally, stimuli in choice probability paradigms targeted a specific feature of the stimulus that a given brain area is known to respond to ([Newsome et al., 1989, Zohary et al., 1994, Britten et al., 1996, Shadlen and Newsome, 2001]). While cited studies made an important breakthrough in our understanding of the function of the cortical activity, we can further improve our understanding of the neural function by studying responses of neural ensembles to complex natural stimuli. In this study, we ask if classification of complex natural images can be read-out from the activity of neural ensembles in the early (V1) and middle-level visual cortex (V4).

The coding function of neural responses in the visual cortex is often described as a set of filters ([Rao and Ballard, 1999]), where the response of each single neuron acts as a filter of the specific feature of a stimulus (e.g., the orientation). In contrast to such rigid coding schemes, animals use sensory information in a flexible way, showing the ability to manipulate the information with temporal delays and to classify stimuli with respect to abstract rules. Is such flexibility an exclusive property of higher brain areas, or can it be observe already in sensory areas? We propose to study responses of neural ensembles to naturalistic stimuli, where the classification task does not rely on filtering properties of single neurons, but instead requires discrimination of abstract classes of stimuli. In particular, the animal visualizes two stimuli consecutively, interleaved with a delay, and decides whether they are the same (“match”) or different (“non-match”). Since the identity of stimuli changes from one trial to another, correct classification cannot rely on encoding of a particular orientation.

The way neural populations encode task-relevant information could either be a by-product of the activity of single units or else have an intrinsic structure that cannot be explained by the response properties of single units ([Elsayed and Cunningham, 2017]). The organism has to generate behavioral decisions in every trial while the trial-to-trial variability of cortical neurons makes the signal conveyed by single neurons unreliable. Summing activities across neurons can, in general, overcome the variability problem, but only in case where the variability is uncorrelated across neurons ([Britten et al., 1992]). This clearly does not hold for local neural ensembles that share a big proportion of their inputs ([Cohen and Kohn, 2011, Smith and Kohn, 2008]). If neurons are correlated, there should be a non-trivial relation between activities of single neurons and the population code. Recent theoretical and modeling work has shown that the variability of neural responses can be reconciled with deterministic population signals ([Koren and Denève, 2017, Boerlin et al., 2013]). This is possible if we assume that task variables (such as the stimulus and the behavioral choice) are represented by the network as a low-dimensional signal, and the coding of such a signal is distributed across a multitude of neurons with redundant coding function. While coding of low-dimensional variables with spiking neural networks has been laid down in theory, the relation between the activity of single neurons and the signal that is effectively transmitted downstream and contributes to behavior, remains elusive in biological networks of behaving animal agents.

Over the last decade, two major insights have been made on the principles of neural coding in cortical networks. The first insight concerns the relation between the similarity of the tuning function of a pair of neurons (signal correlation) and their common variability across trials (noise correlation). If a pair of neurons has similar tuning, positive noise correlations will harm the information transfer. Vice-versa, if a pair of neurons has dissimilar tuning, positive noise correlation can enhance the quantity of the transferred information ([Averbeck et al., 2006]). In order to maximize the information transfer, neurons with similar tuning therefore should have low correlations, while neurons with dissimilar tuning should have stronger correlations. However, in the brain, tens of thousands of neurons activate during a visual discrimination task. Neural signals are effective only if they are transmitted downstream, and the activity that maximizes the information transfer might result in a neural signal that cannot be reliably read-out by a downstream neuron. The second insight on neural coding and dynamics comes from the observation of neural activity in decision-making paradigms. While many neurons activate during a visual task, in experiments, we only observe a tiny fraction of these. Surprisingly, this small subset of neurons appears to be informative about the choice behavior of animal agents. This can be explained with the existence of coding pools, where a (large) number of neurons share a similar coding function. If within the coding pool neurons are more strongly correlated than across pools, the activity of a randomly chosen neuron from the pool has a high probability of being informative about the behavioral variable. In short, if correlations within the pool are stronger than across pools, such a correlation structure allows to infer task variables from the observation of only a small number of neurons ([Shadlen, 1996, Nienborg et al., 2012]). At the first glance, these two ideas imply divergent views on the role of correlations for coding of behavioral variables, since correlations that harm the information transfer are also those that should be the strongest.

In the present work, we study the coding function of the neural population and relate it to correlations. We assume that the computation that the population is performing is discrimination of binary stimulus classes. We compute the coding function of a single neuron relatively to the activity of the network that surrounds the neuron and also takes into account inter-neuron interactions. We apply an optimal linear decoder that gives the upper bound of the transferred information by a linear classifier ([Vapnik and Vapnik, 1998]). We then extract the structure of population responses, given by the weights of decoding model, to study the relation between the structure of population responses and pair-wise dynamics. Our decoding scheme ensures that the relation between the structure and the dynamics is actually relevant for the behavior ([Panzeri et al., 2017]). As we relate the strength of decoding weights to pair-wise dynamics, we find a strong interplay between the two in the microcircuits in V4, but not in V1. We divide neurons in two mutually exclusive coding pools and show that in V4, but not in V1, correlations within pools are stronger than across pools. In general, the relation between the structure of the population model and correlations show many differences between V1 and V4 microcircuits, which might reflect different stages of information processing, from analysis of the visual scene (V1), towards convergent information about binary choice behavior (V4).

## Results

### Neurons in the brain read-out the activity of projecting units

Two adult monkeys *Macaca mulatta* performed a match-to-sample task on visual stimuli. In every trial, the subject visualized two consecutive stimuli, with a delay period in between (Figure 1A). Stimuli were complex naturalistic images in black and white, lasted 300 ms each and were either identical (condition “match”) or else the test stimulus was rotated with respect to the target stimulus (condition “non-match”). The angle of rotation changed randomly across trials and was adjusted on-line to have on average 70 % of correct responses with non-matching stimuli. While all stimuli fell into one of the two classes, “match” and “non-match”, the identity of the stimuli changed from one recording session to another. The task of the animal was to decide about the similarity of the two stimuli by holding a bar for the choice “different” and releasing the bar for the choice “same”. Laminar arrays with 16 recording channels were inserted perpendicularly to the surface of the brain, recording the multi-unit signal across the cortical depth in V1 and V4 visual areas (see methods, Figure 1B). In both areas, neural populations responded with a transient peak in firing rates, followed by tonic firing (Figure 1C).

**Figure 1.**
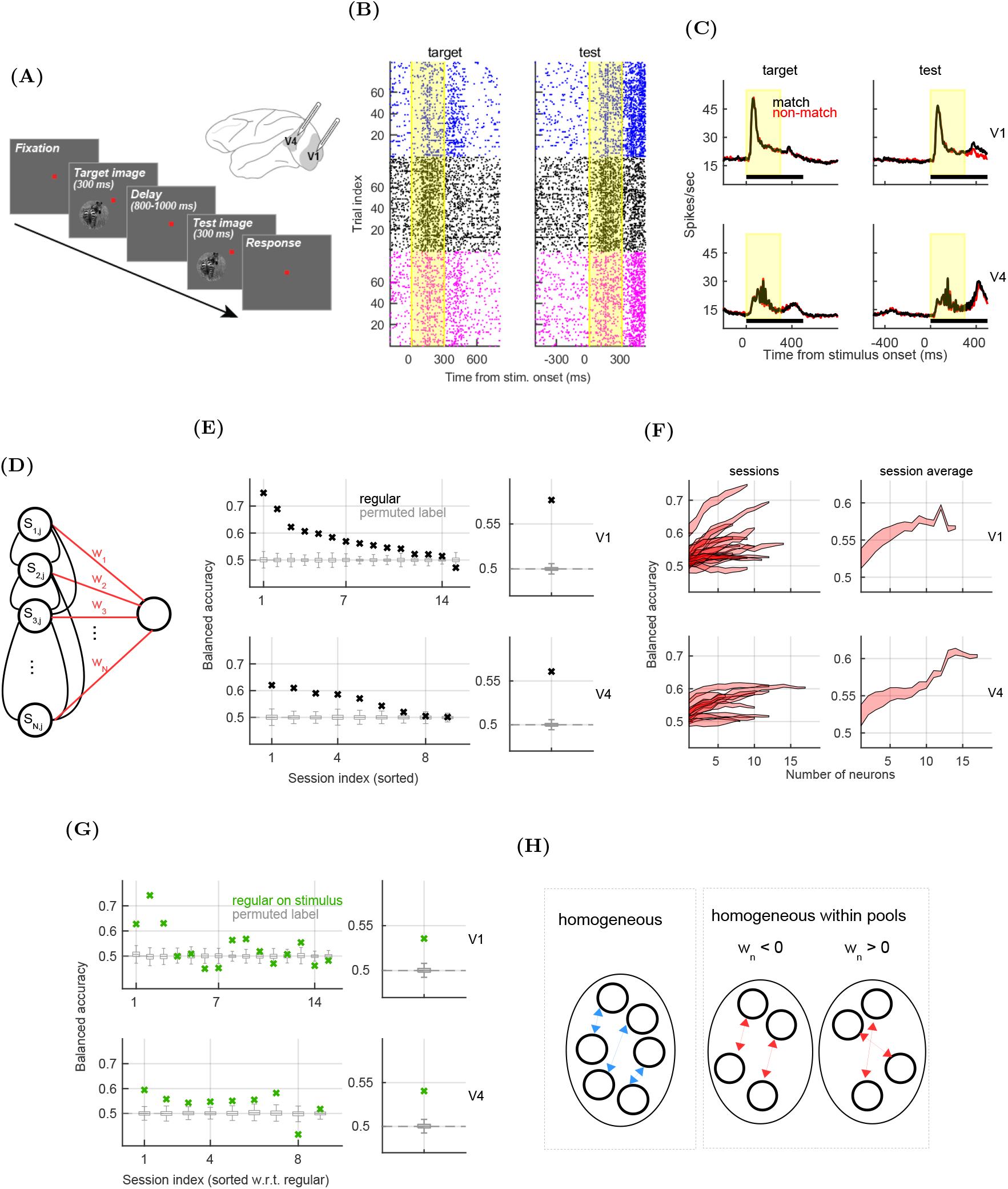
The Population model uses high-dimensional patterns of activation for predicting the stimulus class. **A)** Schema of the experimental paradigm and illustration of the macaque brain, with shaded areas indicating visual areas V1 and V4. **B)** Raster plots of three representative neurons in V4 (blue, black, magenta) in condition “non-match”, during the target (left) and the test time window (right). The yellow area marks the presence of the stimulus. **C)** Population PSTH in V1 (top) and in V4 (bottom), during target (left) and test (right). We separated the activity in conditions “match” (black) and “non-match” (red). **D)** Schema of the population model. Parallel spike counts of the neural population, weighted by the weight vector, project to the read-out neuron. **E)** Left: Prediction accuracy of the population model (black) in V1 (top) and in V4 (bottom). Results have been computed in each recording session and have been sorted, from best to worst performance. Gray boxplots mark distribution of results with permuted class labels. Right: Session-averaged prediction accuracy for the regular model (black) and the model with permutations (gray, *N*_*perm*_ = 1000). **F)** Prediction accuracy in sessions (left) and averaged across sessions (right) as a function of the number of neurons. We plot the mean ± SEM, where the variability is for the random subsampling of neurons, when the number of neurons is less than the total number of neurons in a session (100 subsamples). **G)** Same as in **E**, but for the classification problem with information on the stimulus class and the behavioral performance, but without the information on the behavioral choice. On the left, the order of sessions is the same as in **E**. **H)** Schema of two types of randomization of the structure of parallel spike counts. We permute the neural index across all neurons, without repetition, creating the data that is homogeneous across all neurons (left). On the right, the schema shows permutation of the neural index that is limited to neurons with the same sign of the weight (homogeneous within pools, right).

Neurons and neural ensembles perform their coding function with their dynamics (e.g., by firing more for stimulus 1 than for stimulus 2), and reading-out neural activity with respect to a low-dimensional variable, such as the stimulus class or the behavioral choice, is conceptualized as “decoding”. In decision-making paradigms, decoding of neural activity tells us how accurately one can predict the future behavioral choice of the animal by observing the neural dynamics that precedes it. Here, we investigate decoding models that are readily interpretable in the biological setting, i.e., models that could be implemented in the brain itself. We adopt the view of a read-out neuron that can estimate the change in the firing rate of projecting units in single trials and calculate the probability that such a read-ot neuron would have to discriminate between the two classes of stimuli. We limit our investigation to correct choices, i.e., trials where the visualization of one of the two possible stimulus classes, “match” and “non-match”, has led to the congruent behavioral choice, “same” and “different”. In correct trials, the representation of the stimulus is expected to be the primary source of information for determining the choice behavior. However, at least part of the information about the choice is likely to exist independently of the stimulus, possibly influencing sensory areas as the feedback activity from higher brain areas ([Gilbert and Sigman, 2007]). We therefore assume the existence of a mixed variable, *stimulus* + *choice*, that contains the information about both the stimulus class as well as the behavioral choice (table 1).

**Table 1.** Informational content of task variables.

| Stimulus | Choice | Performance | Info. content |
| --- | --- | --- | --- |
| non-match | “different” | correct | stimulus +<br>choice |
| match | “same” | correct |  |
| non-match | “same” | incorrect | stimulus +<br>performance |
| match | “same” | correct |  |

### Read-out of the activity of populations in V1 and in V4 predicts correct choices better than chance

We decode the variable *stimulus* + *choice* from the activity of neural populations, using an optimal linear classifier, the linear Support Vector Machine (SVM, [Vapnik and Vapnik, 1998]). We call this decoding model the *population model*, since it uses the activation of a population of simultaneously observed neurons (Figure 1D, see methods). The quality of prediction of the population model is assessed on hold out sets with cross-validated balanced accuracy, which is the probability of correctly predicting condition “match” vs. “non-match” in single trials. In case all test samples were classified correctly, we would have the balanced accuracy of 1. The balanced accuracy of 0.5 indicates that the validation samples have been classified equally often correctly as incorrectly, which is the chance level performance. Significance of the prediction accuracy is evaluated with a permutation test, using models with randomly permuted class labels (see methods). It is expected that decoding of the stimulus class should only be better than chance during or after the presentation of the test stimulus, when, potentially, there is information that allows discrimination of “match” from “non-match”. We decoded from the test and the target time window separately, and unless stated otherwise, we used the window [0,500] ms with respect to the onset of the test (target) stimulus. For decoding, we utilized z-scored spike counts, to prevent neurons with stronger firing rate having stronger impact on the decoding model simply because of their stronger firing rate. The range of raw spike counts was bigger in V1 compared to V4 (*p* = 0.001 during test and *p* = 0.006 during target, two-tailed t-test, Figure S1A-B). After z-scoring, the difference of ranges was no longer significant (*p* = 0.692 during test, *p* = 0.325 during target, two-tailed t-test). From the biological perspective, z-scoring corresponds to utilizing the deviation of the spike count from its mean, in units of standard deviation, supposedly more biologically relevant of a signal than the absolute spike count.

Results show that, during the test time window, the population model performs significantly better than chance (*p* < 0.001 in both areas, permutation test, Bonferroni correction, Figure 1D). On average, the high-dimensional model predicts the choice behavior 7.7 % better than chance in V1 and 6.1 % in V4 (Figure 1E). The prediction accuracy varies considerably across recording sessions, presumably because linear arrays are inserted *de novo* in each recording session, capturing small neural ensembles that vary in their relevance for the classification task. As expected, the prediction accuracy increases with the number of neurons (Figure 1F). During the target time window, there is no information available for discriminating matching from non-matching stimuli, and accordingly, the prediction accuracy is at chance (*BAC* = 0.50 in V1 and *BAC* = 0.49 in V4, non-significant, Figure S1C). Interestingly, the use of a nonlinear decoder, SVM with Radial Basis Function, did not improve the prediction accuracy with respect to the linear model (Figure S1D). Since the SVM is the optimal classifier, we conclude that the performance of the population model coincides with the amount of information that a linear read-out on parallel spike counts contains about the variable *stimulus* + *choice*. As we decode from conditions with the information content of *stimulus* + *performance* (conditions that differ in the stimulus class and in the behavioral performance, but do not differ in the choice, table 1), we get the prediction accuracy lower than on *stimulus* + *choice* (3.6 % above chance in V1 and 4 % above chance in V4, Figure 1G).

Prediction accuracy of the population model relies on high-dimensional patterns of activation of the neural ensemble. To get better understanding of this mechanism, we design biologically unrealistic read-out models, that serve the purpose of gaining insights into the population model. As a toy example, consider a network with 2 neurons, where neuron 1 increases its firing rate for the stimulus “non-match” while neuron 2 decreases the firing rate for the same amount, and the opposite happens when the stimulus “match” is shown. A read-out model that is agnostic about the identity of neurons would give chance level prediction, while model that weights contributions of neurons with appropriate weights can utilize the difference in these activation patterns as a source of information. To test this idea on the population model, without changing the dimensionality of the model, we create patterns of neural activity that are homogeneous across neurons. The homogeneous model is given by permuting the neural index of the vector of spike counts in every trial (Figure 1H). Notice that the homogeneous model is equivalent to an unstructured network. As we compare the homogeneous and the regular model, we see that the performance of the homogeneous model is consistently worse (Figure 2A), showing that the patterns of activation of the structured network are an important source of information for discrimination.

**Figure 2.**
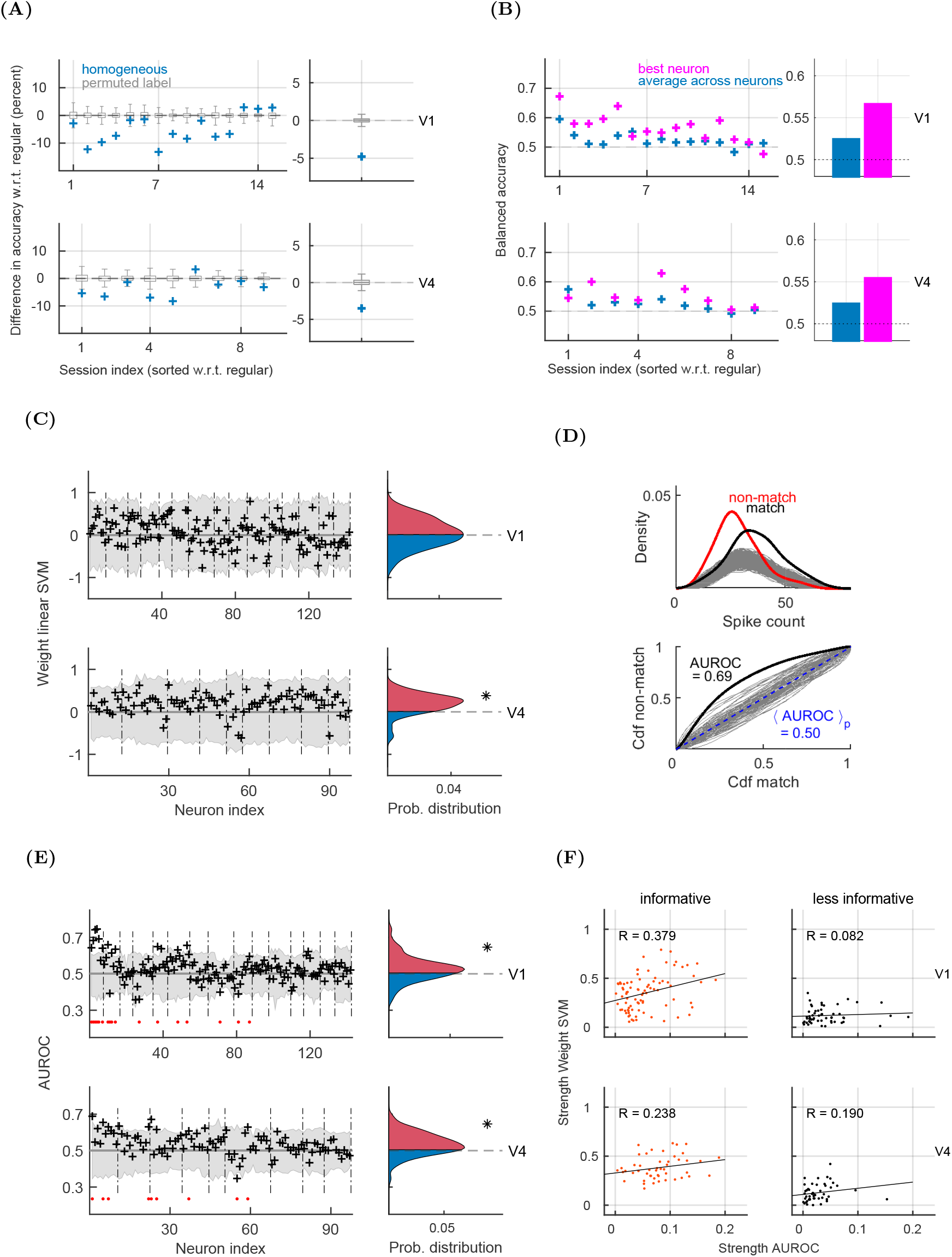
Decoding weights of single neurons are similar, but not identical to population decoding weights. **A)** Difference of the prediction accuracy of the homogeneous model w.r.t. the regular model (homogeneous-regular). Gray boxplots mark the same result for models with permuted class labels. **B)** Left: Prediction accuracy of the best neuron in each session (magenta) and of the uniform pooling (blue). Right: Session-averaged results. The performance of the best neuron has been evaluated with nested cross-validation, using 100 cross-validations for the split of training and test data and 20 cross-validations for the selection of the best model. **C)** Left: Weights of the population model (black), collected across recording sessions in V1 (top) and in V4 (bottom). Dashed vertical lines delimit sessions. The gray area shows the distribution of weights of models with permuted class labels. Right: Distribution of weights, pooled across neurons from all recording session. Asterisk marks significant deviation from 0 (permutation test with Bonferroni correction for the number of neurons in the brain area). **D)** Illustration of the ROC method. The ROC method utilizes probability distributions of spike counts in conditions “non-match” and “match” of a single neuron (top). We calculate the ROC curve by integrating the two distributions with a moving threshold and plotting the two integrals with respect to each other (bottom, black). Gray traces indicate probability distributions (top) and ROC curves (bottom) for the model with randomly permuted class labels for conditions “match” and “non-match”. The blue dashed line is the average across permutations. **E)** Left: Area under the ROC curve (black), with the gray area indicating the distribution of results for models with permuted class labels. Significant neurons are marked with a red dot. Right: Distribution of the Area under the ROC curve, pooled across neurons. The asterisk marks significant deviation from 0. **F)** Scatter plot of the strength of the Area under the ROC versus the strength of weights of the population model, for the subpopulation of informative (left) and less informative neurons (right). *R* marks the Pearson correlation coefficient and the black line is the least squares line. All results are for the test time window and the permutation test is with *N*_*perm*_ = 1000 permutations. Whenever applicable, we plot sessions in the order as on the Figure 1E.

Selecting the best single neuron in each recording session gives performance that is slightly lower than that of the population model. The best single neuron, on average, predicts the *stimulus* + *choice* with accuracy that is 6.6 % better than chance in V1 and 5.4 % in V4 (Figure 2B). The average performance across single neurons, where the prediction accuracy is computed for every neuron independently and then averaged across neurons from the same session (see methods), gives a poor prediction of *stimulus* + *choice*. The prediction accuracy is of 2.4 % above chance in both areas (Figure 2B). While the performance of the best single neuron is, on average, well above the mean performance across single neurons (compare the blue and the magenta bars on the right of the 2B), in some recording sessions, the mean across single neurons gives higher prediction accuracy than the best single neuron (sessions 6 and 15 in V1 and session 1 in V4). This is due to the uncertainty of the estimation of the balanced accuracy.

### Discriminatory power of single neurons is positively correlated with the strength of the weight of the population model

The population model relies on a high-dimensional plane in the space of inputs, fully defined by the vector of weights and the offset of the plane from the origin. Weights describe the role of each neuron for the classification task, relatively to the activity of other neurons from the population, and by taking into account inter-neuron interactions. An identical activity of a single neuron would results in a different decoding weight, if this neuron was surrounded by a different population. If the activity of a particular neuron has no role in discrimination, its weight will be 0. The bigger the deviation of neuron’s weight from 0, the stronger the importance of the neuron for discrimination. The sign of the weight is also important to consider, since weights of the opposite sign have the opposite effect on the direction of the separating hyperplane. Computing weights of the population model for simultaneously recorded neurons during test, and collecting results across recording sessions, we find that almost no neurons have significant weights (Figure 2C, left). Pooling weights across neurons, we find that during test, positive and negative weights are balanced in V1, but unbalanced in V4, with a rather strong excess of positive weights (*p* = 0.005, permutation test, Figure 2C, right). During target, positive and negative weights are balanced in both areas (Figure S2A).

Next, we estimate the discriminatory power of single neurons as the area under the Receiver-Operating characteristic curve (*AUROC*, Figure 2D). The *AUROC* score is the probability of predicting the condition “match” or “non-match” from the distance of distributions of spike counts (see methods). Perfectly overlapping distributions yield *AUROC* = 0.5. Neurons with *AUROC* > 0.5 are those that fire more in condition “match” while neurons with *AUROC* < 0.5 are those that fire more in condition “non-match”. We measure the *AUROC* score for each single neuron and collect results across recording sessions. We find that a small proportion of neurons have *AUROC* significantly different than chance (13 % in V1 and 8 % in V4, 2-tailed permutation test with 1000 permutations, Bonferroni correction, Figure 2E, left). As we pool *AUROC* scores across neurons, we find that during the test time window, there are more neurons that fire stronger in condition “match” compared to “non-match” (*p* < 0.001 in V1 and in V4, permutation test on the sum of *AUROC* scores, Figure 2E, right). The shift of the mean *AUROC* from 0.5 is of 3% in V1 and of 4 % in V4. During the target, the distribution of AUROC scores is centered around 0.5 in both areas (Figure S2B).

Finally, we ask, how do weights of the population model relate to discriminatory capacity of single neurons.

We compare the strength of population weights, 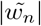, with the strength of *AUROC* scores, |*AUROC*_*n*_ − 0.5|. The strength of the weight is the proxy of the importance of the activity of a particular neuron for the population model. We divide neurons in informative and less informative by ranking the absolute value of the weight of each neuron within the distribution of the same result from models with permuted class labels. The threshold for informativeness is set in such a way as to have the groups of informative and less informative neurons approximately balanced (*α* = 0.3). The strength of the weights and of *AUROC* scores is moderately positively correlated for informative neurons in V1 and for both informative and less informative in V4 (Figure 2F). For less informative neurons in V1, the strength of weights and of *AUROC* is only weakly correlated (Figure 2F, top right). As we distinguish neurons with positive weights (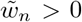, plus neurons) and neurons with negative weights (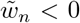, minus neurons), the strength of weights and of *AUROC* scores is moderately (V1) to strongly (V4) positively correlated for both groups (Figure 3A). The weights of the population model are therefore positively correlated with the discriminatory power of single neurons, meaning that the discriminatory capacity of single neurons is an important source of information for the population model. Nevertheless, the fact that the two measures are not identical indicates that the population model has an intrinsic structure that, in particular in V1, cannot be accounted for by the activity of single neurons.

**Figure 3.**
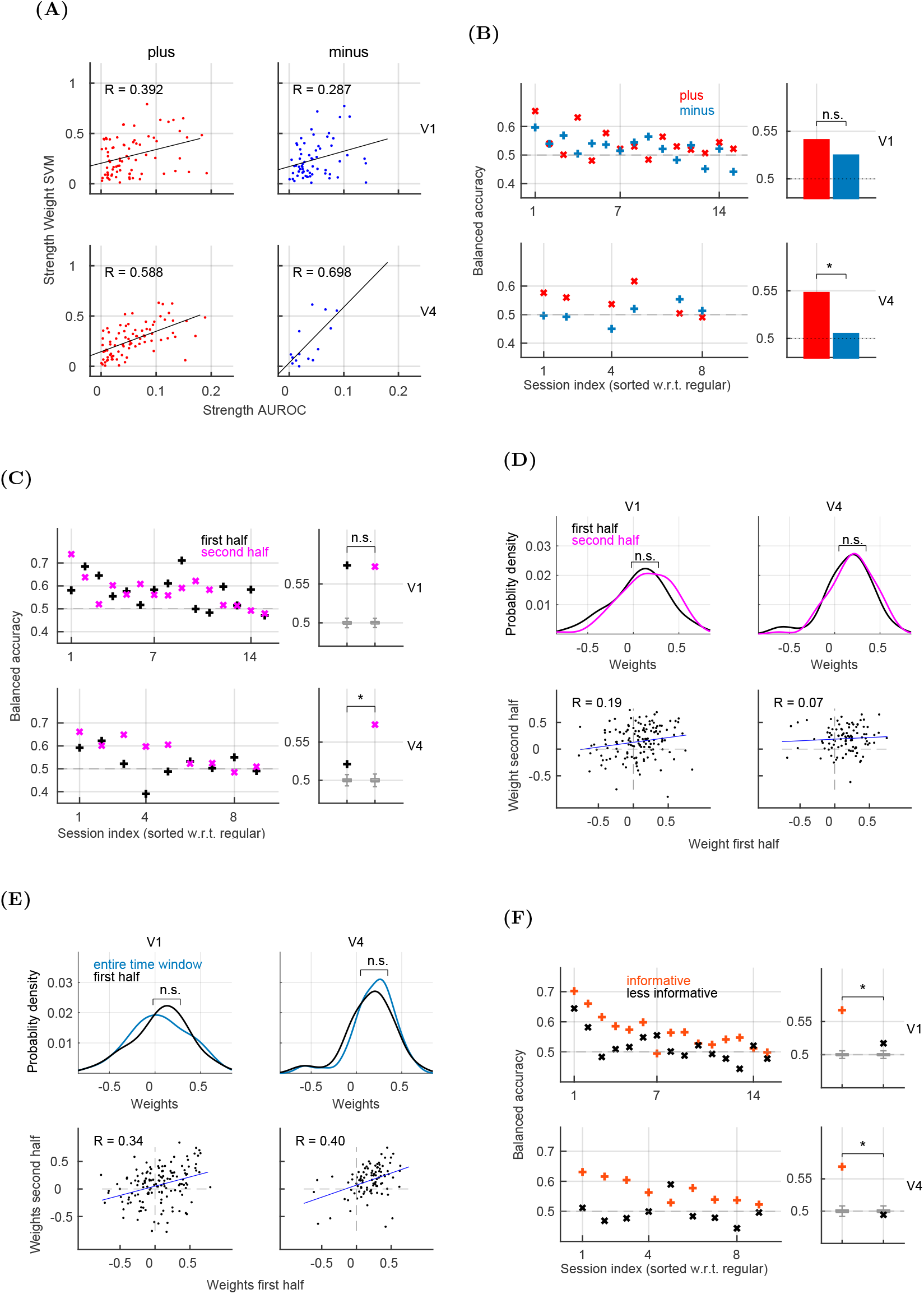
The population model changes between the first and the second half of the test time window. **A)** Scatter plot of the strength of the Area under the ROC versus the strength of weights of the population model, for the subpopulation of neurons with positive weight (left) and negative weight (right). *R* marks the Pearson correlation coefficient and the black line is the least squares line. **B)** Prediction accuracy of neurons with positive (red) and negative (blue) weights. We plot results in recording sessions on the left, and session-averaged results on the right. Asterisk marks significance (permutation test). **C)** Left: Prediction accuracy in recording sessions during the first half (black) and the second half (magenta) of the test time window. Right: Session-averaged results and distribution of results for models with permuted class labels (boxplots). **D)** Top: Distribution of weights for the first (black) and the second half (magenta) of the time window. We plot results in V1 (left) and in V4 (right). Bottom: scatter plot of weights in the first vs. second half of the time window. *R* is a linear correlation coefficient. **E)** Same as in **E**, but comparing weights in the first half of the time window and the weights for the entire time window. **F)** Left: Prediction accuracy for informative neurons (orange) and less informative neurons (black) in V1 (top) and in V4 (bottom). Right: Session-averaged results and the distribution of results for models with permuted class labels (gray boxplots). All results are for the test stimulus and significance is assessed with the permutation test (*N*_*perm*_ = 1000). When applicable, the order of sessions is the same as for the regular model (Figure 1E).

### In V1, prediction accuracy is stable across the trial, while in V4, it increases between the first and the second half of the trial

In the following, we divide neurons with respect to the sign of the weight into plus and minus subpopulation and compute the population model using the activity of each subpopulation. The number of neurons is balanced across groups in each recording session, which requires downsampling the group with more neurons (using *N*_*perm*_ = 1000 random permutations of the neural index) and discarding 3 recording sessions in V4, where there are no neurons with negative weight. With this procedure, we find that plus and minus neurons in V1 and plus neurons in V4 predict the variable *stimulus* + *choce* better than chance (*p* < 0.001 for plus in V1, *p* = 0.005 for minus in V1 and *p* = 0.007 for plus in V4, permutation test), while minus neurons in V4 predict at chance (*p* = 0.414). In V4, the activity of plus neurons is clearly more predictive than the activity of minus neurons (*p* = 0.009, permutation test), while in V1, the difference is not significant (*p* = 0.040, not significant after the Bonferroni correction, Figure 3B). We obtained similar results when, instead of downsampling, we use all neurons from the recording session, but remove the information from a specific group of neurons (Figure S3A-B).

During the trial, neural populations are expected to receive inputs from different sources. In particular, we would expect that neuronal activity is primarily driven by feed-forward inputs at the beginning of the trial, while local and top-down inputs might be important during the second part of the trial. We divide the time window of [0,500] ms w.r.t. the stimulus onset, utilized thus far, into first half ([0,250]ms) and second half of the trial ([250,500]ms w.r.t. stimulus onset). As we compute the prediction accuracy for each window, results show that, on average, the performance is stable in V1, but strongly increases from the first to the second half of the trial in V4 (Figure 3C). Even though the average performance is stable in V1, it differs in individual recording sessions, suggesting different underlying classification models. While the distribution of weights between the first and second part of the trial remains unchanged (Figure 3D, upper plots), the weights of individual neurons change considerably. Between the first and the second half of the trial, weights are only moderately positively correlated in V1 (R=0.19) and almost uncorrelated in V4 (R=0.07, Figure 3D, lower plots). As expected, stronger correlation are observed as we compare the weights from the first (Figure 3E) and second half (Figure S3C) to weights from the entire time window. We remark that, during target, weights between the first and second part of the trial are uncorrelated in both brain areas (Figure S3D).

### In V4, but not in V1, informative neurons are more strongly coupled, synchronized and correlated than less informative neurons

Next, we ask whether the information from neurons with strong weights is sufficient to reproduce the prediction accuracy of the population model. To test this, we keep activation of informative neurons intact, and replace the activation of less informative neurons with the activation from a randomly chosen trial, using the random permutation of the trial index. This way, the group of less informative neurons only contributes noise to the model. Prediction accuracy of such a model in V1 is slightly but significantly lower compared to the regular model (*p* < 0.001, permutation test), and not significantly different from the regular model in V4 (Figure 3F). As we now replace activation of informative neurons with activation from a random trial, and keep activation of less informative neurons intact, we get chance level prediction in V4 and slightly higher than chance prediction in V1. Notice that these results depend on the threshold for splitting neurons in informative and less informative. Nevertheless, they suggest that decoding from the subpopulation of neurons with strong weights, the decoder is almost as good as when decoding from all neurons.

Next, we ask whether informative neurons differ from less informative neurons in the strength of coupling of single neurons to the population (Figure 4A). The coupling function is computed as the strength of the population activity around the spike time of a selected neuron (see methods). Examples of the coupling function are plotted on Figure 4B (left). We find that in V4, informative neurons are more strongly coupled to the rest of the population than less informative neurons (*p* < 0.001, permutation test), while no effect is observed in V1 (*p* = 0.654, Figure 4B, middle and right).

**Figure 4.**
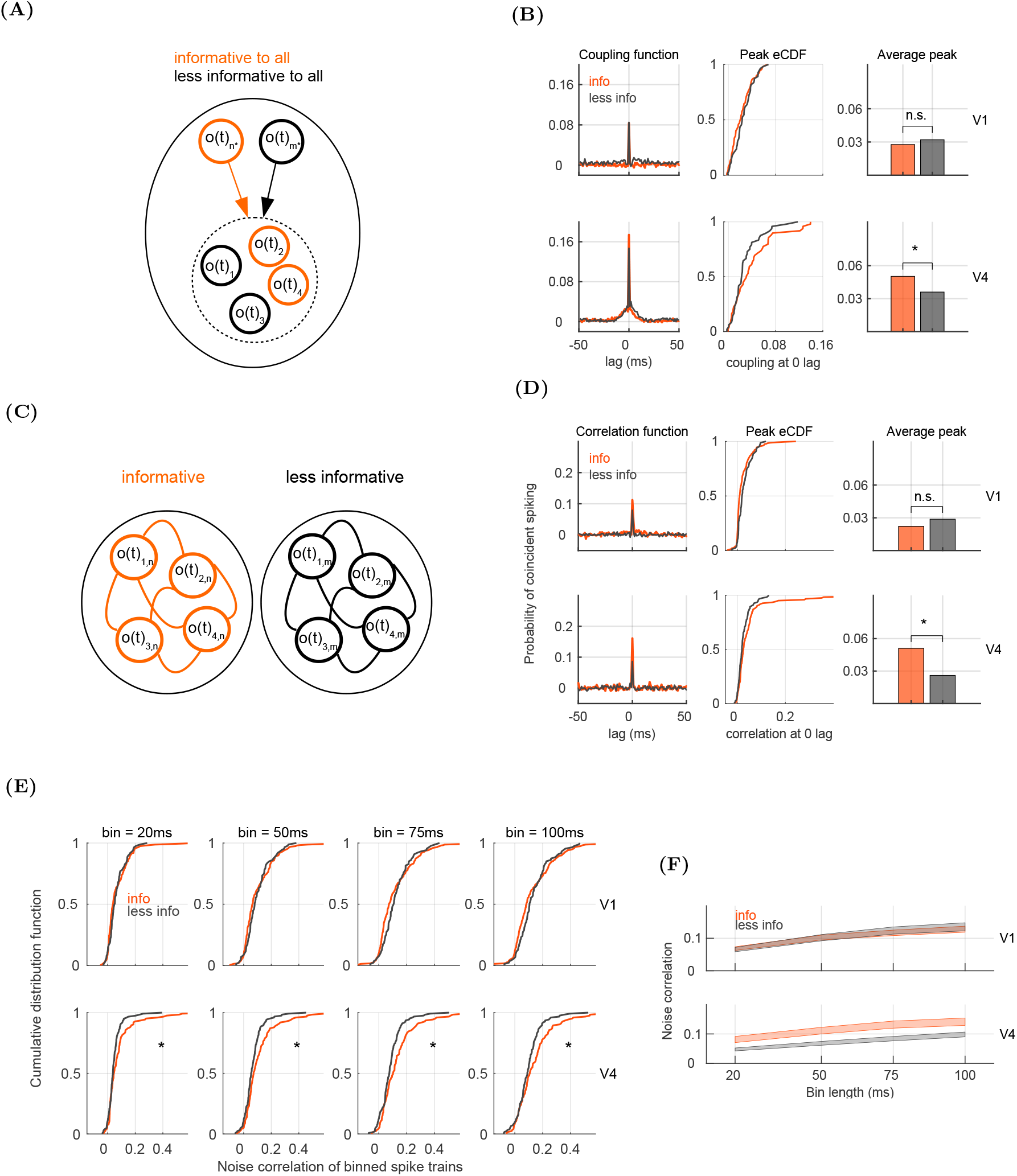
In V4, synchrony and correlations among pairs of informative neurons are stronger than among pairs of less informative neurons. **A)** Schema of coupling of the spiking activity of informative (orange) and less informative (black) neurons to the population activity. The population activity comprises the activity of all neurons but the selected one. **B)** Left: Coupling function for an example informative (orange) and less informative (black) neuron. Middle: Empirical cumulative distribution function (CDF) of the peak of the coupling function at zero lag for informative (orange) and less informative neurons (black). Right: Average peak for the two groups of neurons. Asterisk marks significance (permutation test). **C)** Schema of pair-wise interactions within the group of informative (orange) and less informative (black) neurons. **D)** Left: Correlation function for an example pair of informative and less informative neurons. Middle: Empirical CDF of the correlation function at zero lag (synchrony). Right: Average synchrony for pairs of informative and less informative neurons. **E)** Noise correlations of spike counts for pairs of informative and less informative neurons. We plot the empirical CDF of correlation coefficients, computed for bin sizes of [20,50,75,100] ms. **F)** Mean ± SEM for results in **E**. All results are for the test stimulus and using *N*_*perm*_ = 1000.

Finally, we ask whether informative and less informative neurons differ in synchrony of spiking and correlations of spike counts ([Bair et al., 2001], see methods). Synchrony and correlations are computed for pairs of neurons within the group of informative and less informative neurons (Figure 4C). Representative examples of the correlation function are plotted on the left of the Figure 4D. As we test the height of the correlation function at zero time lag (i.e., pair-wise synchrony) for informative vs. less informative pairs, we see that in V4, synchrony is stronger among informative neurons (Figure 4D, middle and right). Same result is observed as the correlation function is computed during the first half and the second half of the test time window (Figure S4). As we measure noise correlations of binned spike trains, with spike counts in bins of the length of [20, 50, 75, 100] ms, correlations between informative neurons are stronger than between less informative neurons (*p* < 0.001 for bin length of 20, 50 and 75 ms and *p* = 0.003 for bin length of 100 ms, Figure 4E-F). Note that this effect consistently takes place during test in V4, but not in V1, and neither during target in V4 (Figure S5).

### In V4, but not in V1, coupling, correlations and spiking synchrony within the coding pool are stronger than across pools

Besides the strength of the weight, the sign of the weight is important for the structure of population responses, since neurons with the opposite weight have the opposing effect on the angle of the classification boundary (see methods). By dividing the population in neurons with positive and negative weights, we obtain two mutually exclusive coding pools. We showed that removing the structure of population responses across all neurons causes the population model to loose most of its predictive power (Figure 2A). We now ask, how much of the predictive power is lost if we selectively remove the structure only within coding pools (Figure 1H, right). The model that is homogeneous within coding pools performs better than chance in both areas (3.7 % better than chance in V1 and 5.4 % in V4). Comparing the model with homogeneous pools to the regular model, the model with homogeneous pools is significantly worse in V1 (*p* < 0.001, permutation test, Figure 5A), but not in V4, where the difference is not significant (*p* = 0.062). This means that in V4, the information about the sign of the weight is sufficient to reproduce the performance of the regular model, and the information on the amplitude of weights does not add much information. Note that in V4, the size of coding pools is heavily imbalanced, with an excess of plus neurons, and that only plus neurons perform better than chance (Figure 3B). Permutation within the pool of plus neurons must therefore engage a disproportionately big pool of neurons. In spite of that, the prediction accuracy did not drop significantly w.r.t. the regular model. These results persist when we compute the prediction accuracy separately for the first and the second half of the trial (Figure 5B). In V4, the performance of the model with homogeneous pools is similar to the regular model, low during the first half and higher during the second half. In V1, meanwhile, the performance with homogeneous pools is slightly but consistently lower than the regular model in all time windows (compare Figure 5B and Figure 3C), showing again that in V1, the amplitude of weights contributes to classification performance.

**Figure 5.**
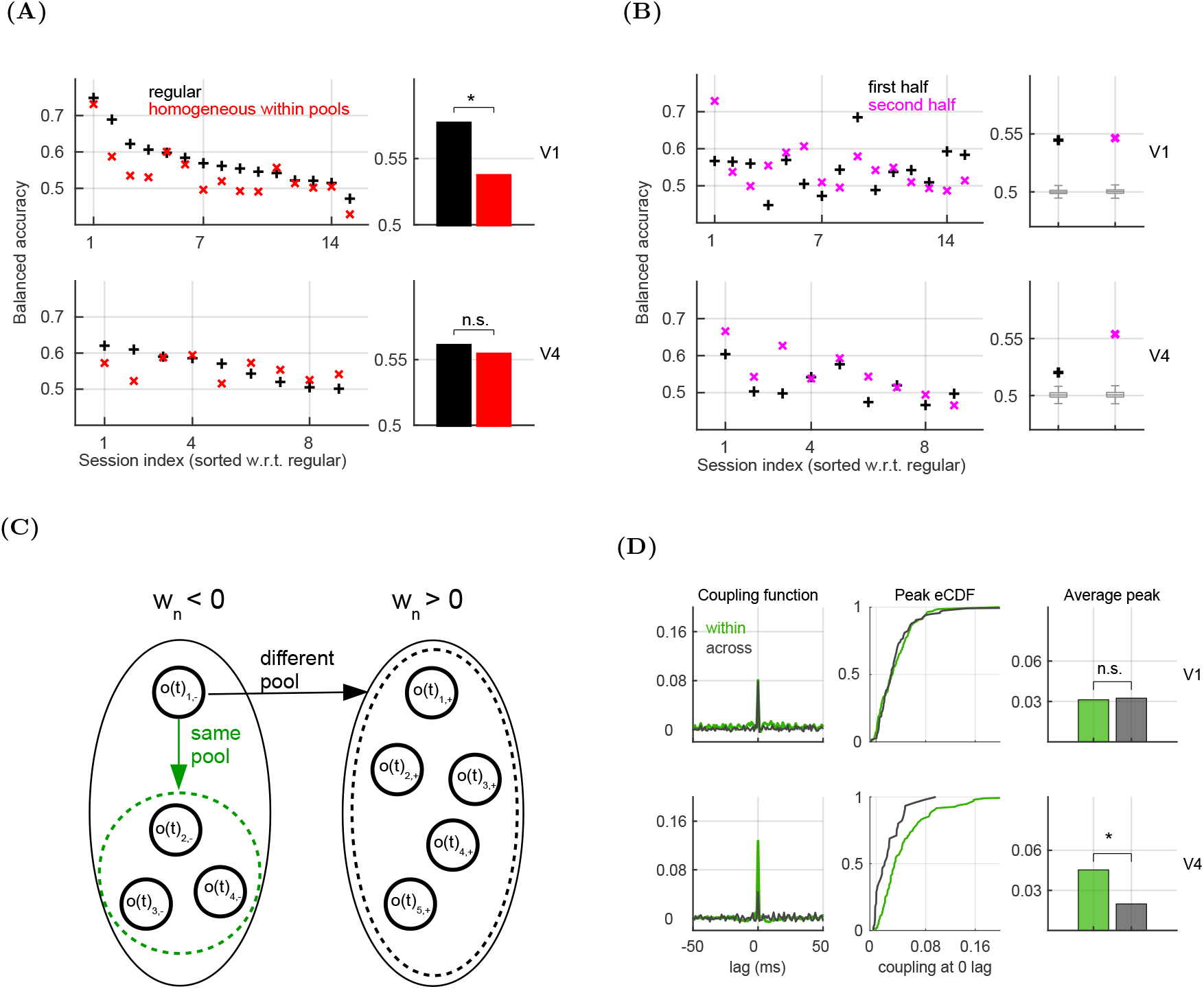
In V4, coupling within pools is stronger than across pools. **A)** Prediction accuracy of the model that is homogeneous across neurons in the same coding pool. The homogeneity of pools is achieved by permuting the neural index among neurons with the same sign of the weight. We use the entire time window ([0,500] ms w.r.t. stimulus onset). The order of sessions is the same as for the regular model (Figure 1E). **B)** Same as in **A**, but using the first half (black) and the second half of the time window (magenta). **C)** Schema of coupling within pool (green arrow) and across pools (black arrow). The dashed line delimits the group of neurons, used to compute the population activity. **D)** Left: The coupling function for an example neuron, for coupling within the pool (green) and across pools (black). The coupling function is plotted as a function of the time lag and is computed in the entire time window ([0,500] ms w.r.t. the stimulus onset). Middle: Empirical CDF for coupling at 0 lag, distinguishing coupling within and across pools. Right: Average coupling at 0 lag, with asterisk marking significance (permutation test). All results are for the test stimulus and the permutation tests utilize *N*_*perm*_ = 1000 permutations.

Next, we compare the strength of coupling of single neurons to the population, distinguishing coupling within pools and across pools. For coupling within pools, the selected neuron and the neural population are from the same coding pool, while for coupling across pools, the selected neuron and the population are in different pools (Figure 5C). During the test time window in V4, coupling is significantly stronger within the pool compared to across pools (*p* < 0.001, Figure 5D), while there is no effect in V1 (*p* = 0.654). Same result is observed as we split the trial in the first half (*p* < 0.001 in V4, *p* = 0.170 in V1) and the second half (*p* < 0.001 in V4, *p* = 0.092 in V1, Figure S6 A-B).

Finally, we compare the strength of pair-wise interactions for pairs within and across coding pools (Figure 6A). We test the spiking synchrony and noise correlations on progressively longer time scales. Noise correlations are measured on spike counts in binned spike trains, for bin length of [20,50,75,100] ms and as noise correlations of the trial-to-trial variability in the 500 ms time window (see methods). Notice that these measures assess pair-wise interactions on the time scale ranging from 1 ms to 500 ms. In V4, pair-wise synchrony is stronger for pairs within compared to across coding pools (*p* = 0.001, Figure 6B). Same effect is observed when we measure the correlation function in the first half and second half of the trial (in V4, *p* < 0.001 for the first half and *p* = 0.001 in the second half, Figure S6 C-D). Consistently, noise correlations of spike counts in V4 are stronger within compared to across pools for all tested bin lengths (*p* = 0.004 for bin length of 20 ms, *p* < 0.001 for 50 ms, *p* = 0.020 for 75 ms, and *p* = 0.032 for 100 ms, Figure 6C-D). Also, noise correlations of trial-to-trial variability are stronger within compared to across pools (Figure 6E). The effect is again only observed in V4 (*p* < 0.001), but not in V1 (*p* = 0.731). In summary, during test in V4, pair-wise interactions on different time scales all reflect the functional separation of neurons into coding pools. This effect is absent in V1, where pair-wise interactions within and across pools are of similar magnitude.

**Figure 6.**
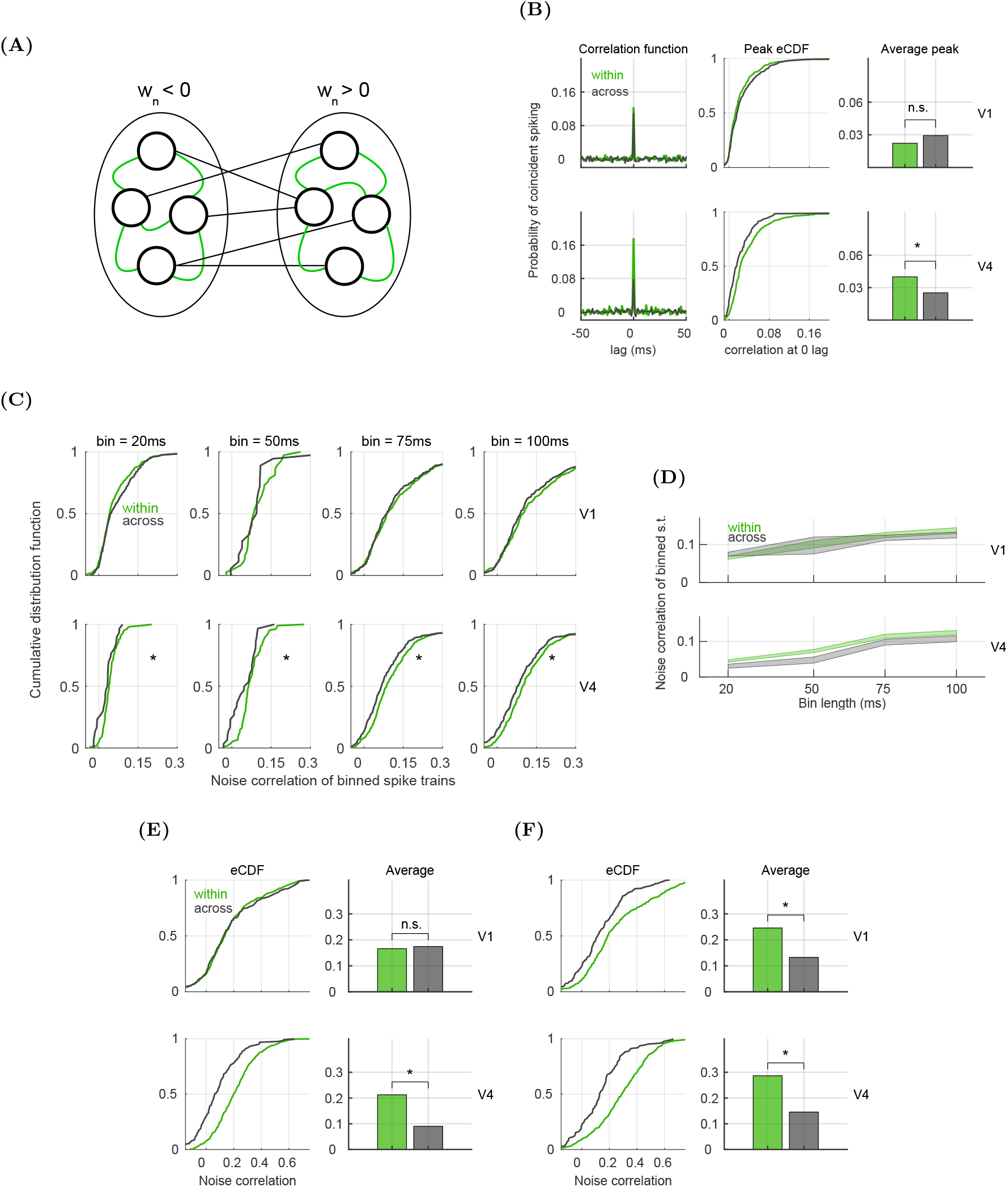
In V4, pair-wise synchrony and correlations within coding pools are stronger than across pools. **A)** Schema of two coding pools, one with negative (left) and the other with positive (right) weights. Green links are for pair-wise interactions within the pool and black are for interactions across pools. **B)** Left: Correlation function for two representative pairs of neurons, one pair from the same coding pool (green), and one pair where neurons are from different pools (black). Middle: Empirical CDF for the correlation function at zero lag (synchrony). Right: Average synchrony within and across coding pools. For all, we use the entire time window ([0,500] ms w.r.t. the stimulus onset). **C)** Noise correlations of binned spike trains, for bins of [20,50,75,100] ms, distinguishing pairs within (green) and across pools (black). We plot the empirical CDF of correlation coefficients, collected across recording sessions, **D)** Same as in **C**, but showing the mean ± SEM of the noise correlations, plotted as a function of the bin length. **E)** Noise correlations of trial-to-trial variability for pairs of neurons within the coding pool (green) and across pools (black). We plot the empirical CDF (left) and the average correlation coefficient (right). For the latter, asterisk marks significance (permutation test). **F)** Same as in **E**, but for the target time window. For all plots, we used the entire time window ([0,500] ms w.r.t. the stimulus onset) and *N*_*perm*_ = 1000 permutations.

Interestingly, during the target time window, correlations of trial-to-trial variability are stronger within compared to across pools not only in V4 but also in V1 (*p* < 0.001 in both areas, Figure 6F). During the first half of the test time window, correlations within pools are stronger than across pools in both areas (*p* < 0.001 in both, Figure S6E). During the second half of the trial, this effect vanishes in V1 while it persists in V4 (*p* = 0.067 in V1, *p* < 0.001 in V4, Figure S6F). These results suggest that stronger correlations within pools might be the default in both V1 and V4, and that the effect vanishes from V1 during the second half of the test time window, possibly to improve the information transfer of the activity of V1 neurons to downstream areas.

### Correlations within pools are harmful for the decoder while correlations across pools are not

In the last part, we measure the effect of correlations on the classification performance of the population model by removing the correlation structure between (a specific group of) neurons and testing whether such a perturbation changes significantly the classification performance with respect to the regular model. The removal of the correlation structure is achieved by permuting the order of trials, independently for each neuron within the group of interest. We perturb the data in two ways: 1.) by removing correlations across the two coding pools but keeping correlations within the coding pools intact and 2.) by removing all correlations (Figure 7A). Note that these two types of correlations are mutually exclusive and comprise all correlations in the network. Removal of correlations across the coding pools does not affect the classification performance (Figure 7B), however, removing all correlations increases the classification performance in both brain areas (*p* < 0.001 in V1 and V4, Figure 7C). The increase in performance occurs mostly during the second half of the trial (Figure 7D), in particular in V4, where the increase is limited to the second half. Testing the effect of the perturbation of the type 1) in the first and second half of the trial, we see no significant effect in V4, and a small increase in performance during the second half in V1 (Figure 7E). As we compare weights of models with removed noise correlations with weights of the regular model, we see that removing noise correlations causes shifting of the distribution towards negative weights (Figure 7F).

**Figure 7.**
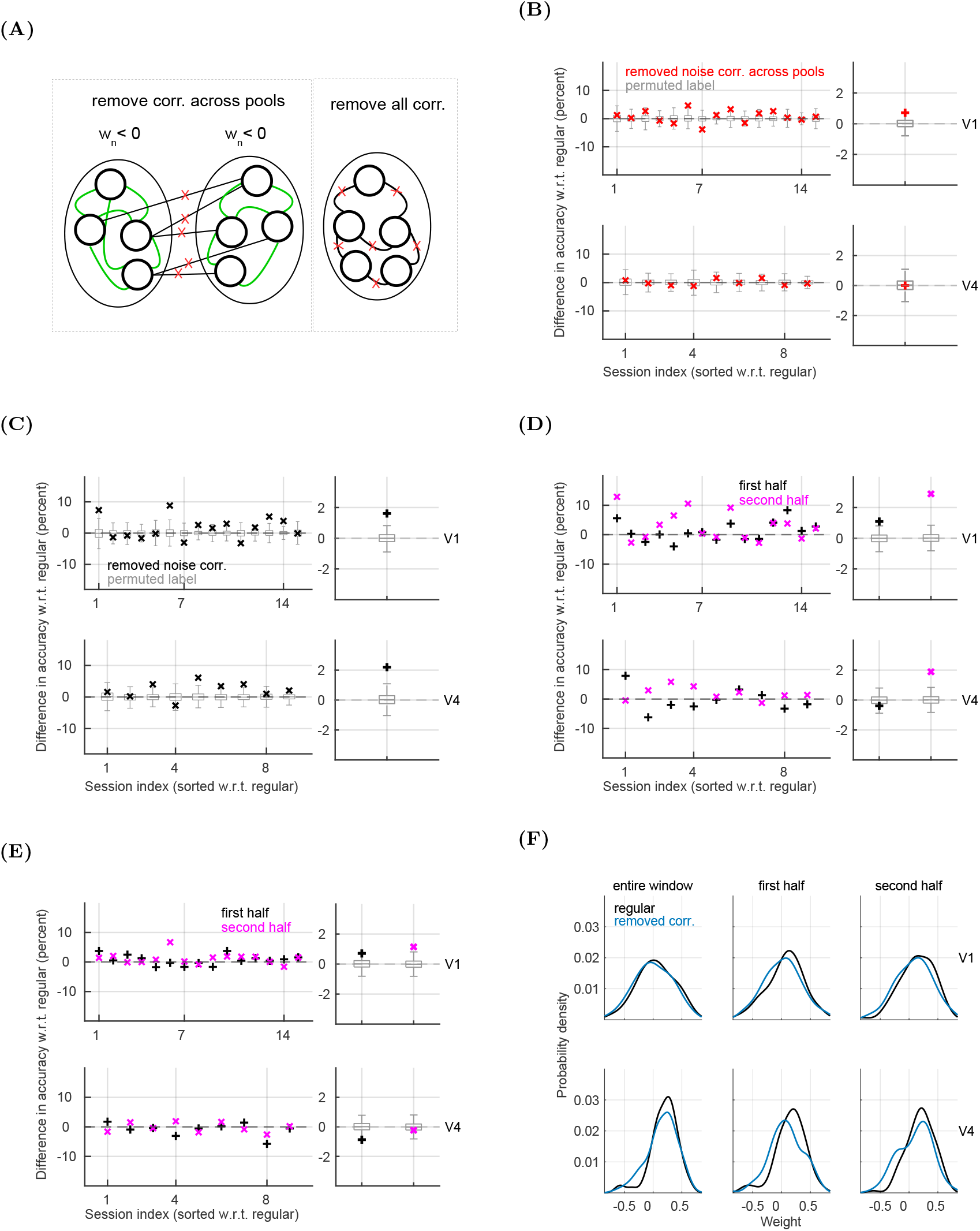
Removing correlations within coding pools increases the predictive power of the population model. **A)** Schema of selectively removing correlations across coding pools, while correlations within pools are kept intact (left), and of removing correlations across all neurons (right). **B)** Difference in prediction accuracy between models with removed correlations across pools and the regular model. We show results in recording sessions (left) and session-averages (right). The entire time window has been utilized. **C)** Same as in **B**, but for the removal of all correlations. **D)** Same as in **C**, but for the first (black) and second (magenta) half of the time window. **E)** Same as in **B**, but for the first (black) and second (magenta) half of the time window. For plots **B** to **E**, the testing is done with the permutation test, using *N*_*perm*_ = 1000 permutations. **F)** Distribution of weights for the regular model (black) and the model with removed noise correlations across all neurons (blue). We plot results for the entire window (left), first half of the window (middle) and second half (right). For all plots, when applicable, we use the permutation test with *N*_*perm*_ = 1000 permutations and order sessions as for the regular model (Figure 1E).

## Discussion

We addressed the structure of parallel spike counts and the role of noise correlations for discrimination of binary variable *stimulus* + *choice* from the activity of neural ensembles in V1 and V4 of the macaque. We showed that the population model predicts the stimulus class with accuracy that is better than chance in both brain areas. The population model utilizes high-dimensional patterns of activation of single neurons as the most important source of information for discrimination. The strength of population weights is positively correlated with discriminatory capacity of single neurons, however, the population model cannot be fully accounted for by aggregating responses of single neurons. In V4, but not in V1, the structure of the population model is clearly reflected in pair-wise dynamics on short and middle-length time scales. We show that the discrimination capacity of the population model relies on a subset of most informative cells, that are in V4 also more strongly coupled, synchronized and correlated than less informative neurons. As the population is split in two coding pools according to the sign of decoding weight, we show that in V4, but not in V1, pairs of neurons within the coding pool are more strongly coupled, synchronized and correlated than across pools. The effect of the coding function on the correlation structure consistently appears on time scales going from 1 to 500 ms. Finally, we show that correlations within the pool are harmful for the decoder, while correlation across pools are not.

While the choice probability has already been reported in V4 ([Shiozaki et al., 2012]) as well as in V1 ([Nienborg and Cumming, 2014]), previous studies used artificial stimuli that specifically target filtering properties of observed neurons. While artificial stimuli might potentially maximize the choice probability in observed neurons, they are also likely to affect inter-neuron correlations. We argue that our study adds an important piece of information to existing literature on choice probabilities by introducing naturalistic stimuli, and a delayed-to-sample visual task. The present behavioral task requires to memorize the template image (the target stimulus) and compare it to a delayed sample (the test stimulus), and is expected to activate not only multiple brain areas across the visual ventral stream, but also the prefrontal cortex ([Gold and Shadlen, 2007]). Moreover, the orientation of the test stimulus changes from one trial to another, which prevents the discrimination to rely on the specific orientation of the test stimulus. In such a setting, it is not trivial to predict the stimulus class in early (V1) and middle (V4) visual cortex with probability that is better than chance.

Our results suggest that the predictability of the stimulus class could be due to the local structure of neural ensembles and/or to top-down projections from higher brain areas, such as IT or prefrontal cortex. The classification of matching and non-matching stimuli could be computed downstream, and the choice-related signal would then be projected back to V1 and V4 to reinforce the internal evidence for the choice. All other inputs being equal, the population model might be capturing a difference in top-down projections for matching and non-matching stimuli. Local separation into coding pools is additionally supported by the correlation structure, where neurons within the coding pool are more strongly correlated than neurons across pools. Present results suggest that having stronger correlation within compared to across pools might be the default correlation structure in the present experimental task, that vanishes from V1 responses towards the end of the test stimulus. Since correlations within pools are harmful for the prediction accuracy, correlations during test in V1 might be actively suppressed in order to improve the information transfer of the choice-related signal. Since pairwise correlations are either neutral or harmful for the quantity of decoded information, the benefit of high-dimensionality for the information transfer that we show in the present work is in the ability of forming high-dimensional patterns of activity, and not in the correlation structure. However, the fact that informative neurons in V4 have stronger correlations than less informative neurons suggests that correlated activity of informative neurons might be useful for the read-out, perhaps in the sense of making it more reliable. Importantly, related work has shown that coordination of spiking of three and more neurons carries information about perceptual accuracy [Shahidi et al., 2019] and that synergistic interactions of multiple cells across the cortical column are beneficial for the information transfer [Nigam et al., 2019].

Neural microcircuits under investigation present a mixture of correlations, some of which are harmful for the decoder, and others that are not ([Moreno-Bote et al., 2014]). The reason for this might be to perform reasonably well in discrimination tasks, given the constraint of the noisy environment in which neurons operate ([Shadlen and Newsome, 1998]). We argue that in the present experimental task, the activity of neural ensembles has two main functions, one is to collect the information about the stimuli and classify them in “match” and “non-match”, and the other is to calculate a binary variable than can guide the choice behavior. While the neural activity in V1 might be optimized in such a way as to maximize the transfer of the information about the visual scene, the quantity of transferred information might not need to be optimized in downstream areas, where the reliable transmission through correlated channels might be a priority. The repertoire of the choice behavior in the present setting is reduced to a binary variable (the choice “same” and “different”), which only contains 1 bit of information. As correlations with the pool are stronger than across pools, their effect is to amplify the signal within the pool, which might help a reliable transmission of the signal. While most decoding models pose that the read-out unit is a single cell that pools activity from all observed units, recent work has suggested that the pooling of the read-out of the V1 activity is limited to neurons with roughly similar orientation preference (up to 45° difference in the orientation preference, [Andrei et al., 2019]). Taking the selectivity of the read-out into account, each coding pool could be read-out by a distinct read-out unit, thus effectively creating segregated signals from neuronal populations with highly overlapping receptive fields.

If population weights have been learned through a form of Hebbian synaptic plasticity ([Abbott and Nelson, 2000]), the formation of coding pools can be explained by the choice-specific top-down input. On a coarser spatial and temporal scale, it has been demonstrated that patterns of spontaneous activity reflect the functional architecture of the cortex ([Arieli et al. 1996, Tsodyks et al., 1999, Fox and Raichle, 2007]), including the representation of abstract information, such as rules or goals ([Waskom and Wagner, 2017]). Finding the same organization in V4 with the present classification task opens up the possibility that the correlation structure actively supports processing of stimuli in categorization tasks on a much smaller spatial scale.

Prediction accuracy of the population model is the best prediction that can be obtained from a linear classification model in the present classification task, setting the higher bound on the information about the choice that can be extracted from spike counts of observed neural ensembles in V1 and in V4. We showed that the use of a nonlinear SVM with radial basis function kernel does not improve the prediction accuracy. We also argue that the relatively low prediction accuracy of the population model might be importantly amplified as the coding function is exploited dynamically. If population weights are applied on spike trains, weighting the contribution of spikes in real time, the difference between the signals in conditions “match” and “non-match” can accumulate over time. In a related work, we show that a leaky integration of weighted spike trains in conditions “match” and “non-match” results in signals that are significantly different and diverge over the time course of the trial ([Koren et al., 2019]).

It has to be pointed out that while our population model relies on a linear classifier, this linearity should not be understood as the linearity between responses of single neurons and the population. On the contrary, we show that single neurons cannot account for the structure of population responses, given by the weights of the population model. In the biological setting, population weights have a straightforward interpretation as synaptic weights between a population of projecting neurons and a read-out neuron. The sign of the weight would correspond to strengthening (positive weight) or weakening (negative weight) of the synaptic weight between the projecting neuron and the read-out neuron with respect to the baseline. In biological networks, the contributions of projecting neurons sum at the synapse of the read-out neuron and form the synaptic input. What the linearity of the population model does imply is that synaptic inputs sum linearly as they create the synaptic current at the read-out neuron, a well-established experimental fact, and common use in modeling ([Dayan and Abbott, 2001]). While the transfer function of the read-out neuron might not be linear due to the spike-triggered adaptation, voltage-gated currents, etc., this can be easily considered while modeling present results, but is beyond the scope of the present work.

An interesting venue for future research could be the comparison of the orientation tuning of single neurons and their decoding weights. Theoretical studies predict the relation between the similarity of tuning curves and noise correlations ([Averbeck et al., 2006, Moreno-Bote et al., 2014]). In this study, we have related noise correlations and similarity of decoding weights, obtaining similar relation as predicted for tuning curves. In the present experimental setting, computation of the choice-related signal might not directly depend on the orientation tuning of V1 and V4 neurons, and it remains unclear how orientation tuning and decoding weights are related. Future work could also address the variability of weights and how they change over the trial, topics only coarsely studied in the present work. Finally, the separation of feed-forward, local, and feed-back inputs could provide major insights into how neural ensembles in the visual cortex compute choice-related signals.

## Acknowledgments

We would like to thank Henning Sprekeler, Caroline Matthis, Tiziano D’Albis and Youssef Kashef for constructive input on the previous draft.

## STAR Methods

### Lead contact and materials availability

Further information and requests for resources should be directed to and will be fulfilled by Lead Contacts, Veronika Koren and Valentin Dragoi. This study did not generate new unique reagents.

### Experimental model and subject details

#### Ethics statement

All experiments performed in this study were conducted in accordance with protocols approved by The Animal Welfare Committee (AWC) and the Institutional Animal Care and Use Committee (IACUC) for McGovern Medical School at The University of Texas Health Science Center at Houston (UTHealth), and met or exceeded the standards proposed by the National Institutes of Healths Guide for the Care and Use of Laboratory Animals.

#### Animal subjects

Two male rhesus macaques (Macaca mulatta; M1, 7 years old, 15kg; M2, 11 years old, 13kg) were used in this study. Subjects were housed individually (after failed attempts to pair house) in cages sized 73 × 69 × 31 or 73 × 34.5 ×31 inches, in close proximity to monkeys in adjacent cages, allowing for visual, olfactory and auditory contact. Toys were given in rotation, along with various puzzles, movies and radio programming as environmental enrichments. Monkeys were fed a standard monkey biscuit diet (LabDiet), that was supplemented daily with a variety of fruits and vegetables. Subjects had been previously trained to perform visual discrimination task, and each implanted with a titanium head post device and two 19mm recording chambers (Crist Instruments) over V1 and V4. All surgeries were performed aseptically, under general anesthesia maintained and monitored by the veterinary staff from the Center for Laboratory Animal Medicine and Care (CLAMC), with appropriate analgesics as directed by the specialized non-human primate veterinarian at CLAMC. During the study the animals had unrestricted access to fluid, except on days when behavioral tasks were performed. These days, animals had unlimited access to fluid during the behavioral task, receiving fluid for each correctly completed trial. Following the behavioral task, animals were returned to their home cage and were given additional access to fluid. The minimal daily fluid allotment was 50ml/kg (monkeys were weighed weekly), though monkeys could drink more through their participation in the task. During the study, the animals health and welfare was monitored daily by the veterinarians and the animal facility staff at CLAMC and the labs scientists, all specialized with working with non-human primates.

### Method details

#### Experimental procedure

The trial started after 300 ms of successful fixation within the fixation area and consisted in visualizing the target and the test stimuli, with a delay period in between. Stimuli were shown for 300 ms each, while delay period had a random duration between 800 and 1000 ms, in order to prevent the expectation of the timing of the test stimulus. The target and the test stimuli, naturalistic images in black and white, were either identical (condition “match”) or else the test stimulus was rotated with respect to the target stimulus (condition “non-match”). The task of the animal was to decide about the similarity/difference of the target and the test stimuli by holding a bar for “different” and releasing the bar for “same”. The subject had to respond within 200 and 1200 ms from the offset of the test stimulus, otherwise the trial was discarded. The difficulty of the task was calibrated on-line, in such away as to have, on average, 70 percent of correct responses on non-matching stimuli. The difference in orientation of the test stimulus ranged between 3 and 10 degrees.

#### Recordings

In every recording session, two laminar electrodes with 16 recording channels each were inserted in V1 and V4 areas, measuring the multi-unit signal with 16 recording channels (0.1 mm spacing between adjacent contacts). Electrodes were inserted perpendicularly to the cortical surface, and calibrated such that neurons from the two areas had overlapping receptive fields. Since recording sessions were performed on different days, it is extremely unlikely that the electrode captured same neurons in different recording sessions. We therefore treated spike-sorted units from different recording sessions as distinct cells. The analysis included all cells that responded to the stimulus with a 4-fold increase of the firing rate with respect to the baseline. Since we are interested in decoding population activities, we used recording sessions with at least 6 simultaneously recorded units. This amounts to 142 neurons in V1, collected in 15 recording sessions, and to 97 neurons in V4, collected in 9 recording sessions. The average, minimum and maximum number of trials per session are reported in the table 2. Besides mentioned criteria, we did not exclude any part of the data. All the analysis is done for each session individually, and results are then collected across sessions and across the two animal subjects.

**Table 2.** Statistics of trial number per condition.

| area | condition | average trials | min. trials | max. trials |
| --- | --- | --- | --- | --- |
| V1 | “non-match” | 108 | 42 | 230 |
|  | “match” | 118 | 51 | 290 |
| V4 | “non-match” | 73 | 39 | 104 |
|  | “match” | 96 | 78 | 123 |

### Quantification and statistical analysis

Classification and statistical analysis were done with Matlab, Mathworks, version R2017b. All methods were applied to the test and target time windows independently. If not mentioned otherwise, we used the window [0,500] ms with respect to the onset of the corresponding stimulus. For part of the analysis, we used, in addition the the whole window, also the first and second half of the window ([0,250]ms and [250,500]ms w.r.t. the stimulus onset), as mentioned in the main text and in corresponding figures.

#### Classification and extraction of decoding weights

The spike train of a single neuron is a binary vector of zeros and ones,

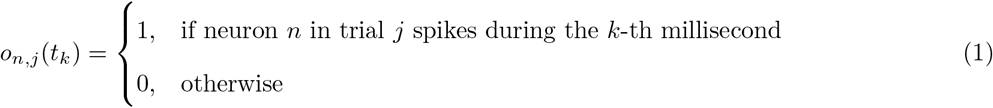

where *n* = 1, …, *N* is the neural index, *j* = 1, %, *J* is the trial index and *k* = 1, …, *K* is the time index with step of 1 millisecond. The 1 millisecond bin is the temporal resolution of the spike train.

#### Area under the ROC curve for univariate analysis

The discriminability of conditions “match” and “non-match” from spike counts of single neurons was evaluated using the receiver operating characteristic (ROC) analysis ([Gold and Shadlen, 2007, Britten et al., 1996, Newsome et al., 1989, Shadlen and Newsome, 2001] [Zohary et al., 1994]). The ROC is a non-parametric method that evaluates the distance between distributions of spike counts in conditions “match” and “non-match”, collected in multiple trials. The spike count of the neuron *n* in trial *j* is defined as the sum of spikes in the desired time window: 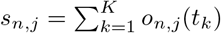.

Spike counts are collected across trials and define two probability distributions, corresponding to conditions “match”, *P*_*M*_(*s*), and “non-match”, *P*_*NM*_(*s*), for *s* ∈ {0, Δ*s*, %, *s*_*max*_}, where *s*_*max*_ is the maximal spike count across all recorded activities, Δ*s* = 2*s*_*max*_/(*nstep* − 1), *nstep* = 200. Probability distributions are convolved with a normalized Gaussian kernel,

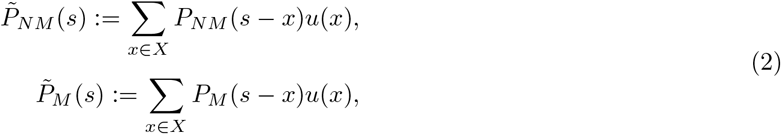

with 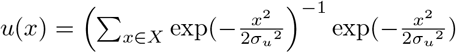. The width of the kernel *σ*_*u*_ is defined as follows: first, an estimate for the optimal width of each distribution is computed using Silverman’s rule, 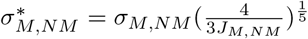, where *σ*_*M*_ (*σ*_*NM*_) is the standard deviation of the distribution of spike counts in condition “match” (“non-match”) and *J*_*M*_(*J*_*NM*_) is the number of trials in condition “match” (“non-match”). Then, we set 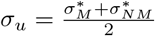.

We define empirical cumulative distribution functions, *D*_*M*_(*s*) and *D*_*NM*_(*s*), as follows:

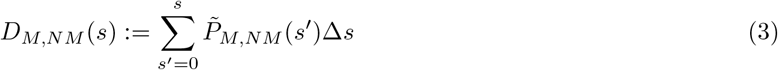

Note that the cumulative distribution functions are bounded, 0 ≤ *D*_*M,NM*_(*s*) ≤ 1, and the area under the ROC curve is the following sum:

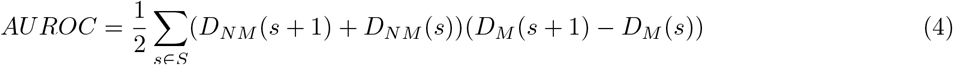

Two identical distributions give the *AUROC* of 0.5, at chance prediction of conditions “match” and “non-match” from the observation of spike counts. In order to estimate if the prediction is significantly better than chance, we have to take into account the variance of the data. This is done with the permutation test. We pool the input statistics (*s*_*n,j*_) from the two conditions and randomly assign labels “match” and “non-match” to every trial. The pooled distribution is then split into two subsets, with the number of samples in subsets corresponding to the number of original samples in conditions “match” and “non-match”. Using these two random subsamples, the area under the curve is then computed by eq. 2 - 4. This procedure is repeated *nperm* = 1000 times, and results in a distribution of areas under the ROC curve, *AUROC*_*p*_, *p* = 1, …, *n*_*perm*_. We compute the fraction of *AUROC*_*p*_ that are larger (*p*_*L*_) and smaller (*p*_*S*_) than the true *AUROC*,

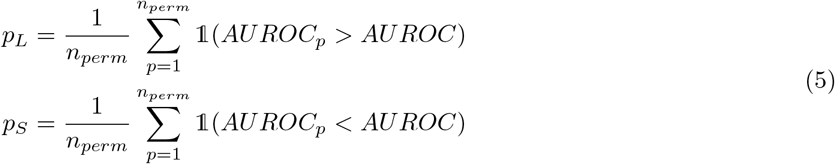

where 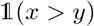 is the following logical function:

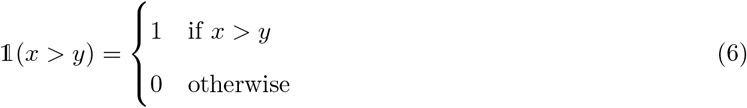

The *AUROC* is considered to be significantly smaller (larger) than 0.5 if the following is true: *p*_*L*_ ≤ ^0.05^/*N*_*tot*_(*p*_*S*_ ≥ ^0.05^/*N*_*tot*_), where *N*_*tot*_ is the total number of neurons in a given brain area.

Finally, we are interested in the distribution of *AUROC* across neurons. We pool *AUROC* across neurons from all recording sessions and from the two animal subjects and get the following distribution: *AUROC*_*n*_, *n* = 1, …, *N*_*tot*_. We compute the mean of the distribution, ⟨*AUROC*⟩_*n*_, and test its significance by comparing it with the distribution of averages from the permutation procedure, ⟨*AUROC*_*p*_⟩_*n*_, *p* = 1, …, *n*_*perm*_. The p-value is then computed by eq. 5-6, and corrected for multiple testing with Bonferroni correction.

#### Linear Support Vector Machine for multivariate analysis

The second read-out method consists in discriminating conditions “match” and “non-match” on high-dimensional activity profiles with linear Support vector machine (SVM). We use z-scored spike counts,

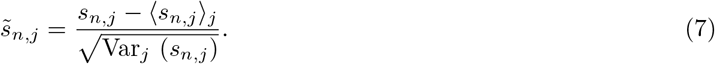

where ⟨*s*_*n,j*_⟩_*j*_ is the empirical mean and Var_*j*_(*s*_*n,j*_) is the empirical variance across trials. One sample for the classifier is an N-dimensional activity vector of N simultaneously recorded neurons in trial *j*,

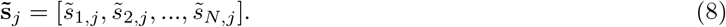

The optimization problem of the linear SVM, in its primal form, is expressed with a Lagrangian [Vapnik and Vapnik, 1998],

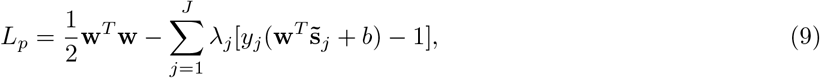

where **w** is (*N,* 1)-dimensional vector of weights and N is the number of simultaneously active neurons, *b* is the offset of the separating hyperplane from the origin, *λ*_*j*_ is the Lagrange multiplier (*λ*_*j*_ ≤ 0 ∀*j*) and *y*_*j*_ is the class label in trial *j*, with *y*_*j*_ ∈ {−1, 1}.

The model is trained on 80 % of trials, and tested on the remaining 20 %. We perform Monte-Carlo cross-validation for random splits in the training and test set with 100 iterations. The classification performance on the test set is computed with balanced accuracy,

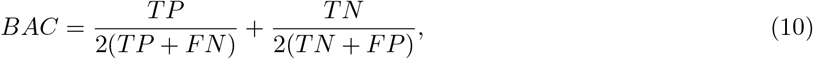

where *TP*, *FN*, *TN* and *FP* stand for the number of true positive, false negative, true negative and false positive classifications on the test set, respectively. We call a true positive the correct classification of condition “match”, a true negative the correct classification of condition “non-match”, etc. Reported balanced accuracy is averaged across cross-validations. The regularization parameter of the SVM is estimated with 10-fold cross-validation on the training data, and chosen from the following range: *C* ∈ {0.0012, 0.00135, 0.0015, 0.002, 0.005, 0.01, 0.05, 0.1, 0.5}.

If the classifier performs at chance, its balanced accuracy is 0.5. The significance of the deviation of balanced accuracy from 0.5 is determined with permutation test. We train models with randomly permuted class labels for “match” and “non-match”. Iterating the permutation procedure gives a distribution of classification performances, *BAC*_*p*_, *p* = 1, …, *n*_*perm*_, with *n*_*perm*_ = 1000. The *p*-value of the permutation test is the probability of models with permutation of having higher balanced accuracy than the true model.

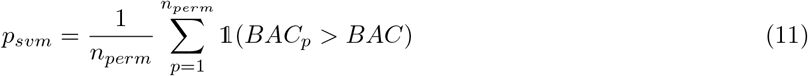

If 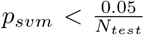, the balanced accuracy is considered to be significantly higher than chance, where *N*_*test*_ = 4 stands for the Bonferroni correction for multiple testing.

We also compute the SVM on single neuron data. The classification procedure is done for each neuron independently. We estimate two measures, the average performance of single neurons and the performance of the best single neuron. For the average performance of single neurons, we compute the balanced accuracy of every single neuron, and average the balanced accuracy across neurons from the same recording session. For evaluating the performance of the best single neuron, we perform nested cross-validation, with the selection of the best neuron in the inner loop, and with the evaluation of the performance of the best neuron in the outer loop. In the outer loop, we re-train the model of the selected neuron using all trials from the inner loop, and test the model on a hold-out set of trials that have not been used in the inner loop. For the split into the training and the test set, the inner loop uses 100 Monte-Carlo cross-validations and the outer loop uses 20 Monte-Carlo cross-validations, and we report averages across cross-validations. We use 80 % of trials for the inner loop and the remaining 20 % for the test set of the outer loop. Note that, because of the limited amount of data and limited number of splits into the training and the test set, all estimates of balanced accuracy come with an uncertainty.

#### Extraction of decoding weights

The minimization of the Lagrangian (eq. 9) allows to compute the vector of weights,

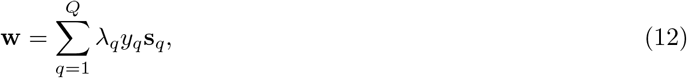

where *Q* is the number of support vectors [Vapnik and Vapnik, 1998]. The range of the weight vector depends on the regularization parameter of the SVM (*C* parameter) and is therefore different from one session to another. Since we combine results from different recording sessions, we normalize the weight vector with *L*^2^ norm,

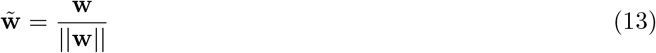

with 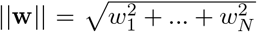. The vector of decoding weights, 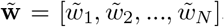, associates the activity of each neuron with it’s role for classification in the *N*-dimensional space of inputs.

The distance of weights is defined as the Euclidean distance of weights between pairs of neurons from the same recording session,

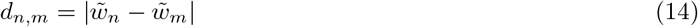

with *n*, *m* = 1, …, *N*. Results are then pooled across recording sessions.

#### Coupling of single neurons with the population

We measure the strength of the coupling of single neurons with the population using a measure of spike-triggered population activity, defined similarly as in methods of Part 1 (see also [Koren and Denève, 2017, Luczak et al., 2013]). We consider the spike train of neuron *n*\* in trial *j*. The spike train can be expressed as a sum of Kronecker delta functions,

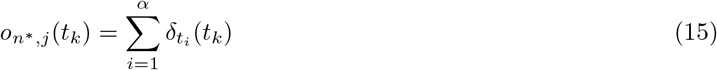

where *t*_*i*_ is the spike time of the *i* − *th* spike and *α* is the number of spikes of neuron *n* in trial *j*. To compute the population activity in trial *j*, we average spike trains across neurons, excluding the neuron *n*\*:

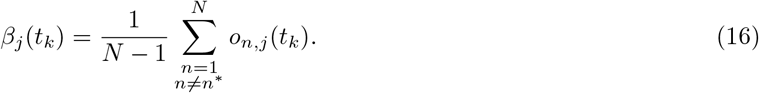

For each spike of neuron *n*\*, we measure the population activity as a function of the time lag before and after the spike:

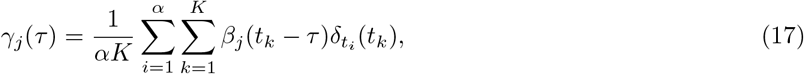

where the lag is *τ* = −*τ*_*max*_, −*τ*_*max*_ + 1, …, *τ*_*max*_. Note that *τ* = 0 corresponds to coupling at the time of the spike. Coupling function *γ*_*j*_(*τ*) depends on the stimulus, and we make it independent by subtracting the trial-invariant coupling. Trial-invariant coupling is computed as coupling where the spike train and the population activity are not from the same trial, e.g., we use 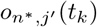 and *β*_*j*_(*t*_*k*_), with *j*′ ≠ *j*. The trial index for the former is determined with random permutation of the order of trials. The permutation is repeated *nperm* = 100 times and the coupling function is averaged across repetitions, resulting in 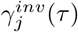. Finally, we obtain the trial-specific (or “noise”) coupling function by subtracting the trial-invariant from the original coupling function, and averaging across trials,

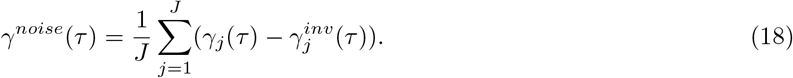

The coupling coefficient is defined as the coupling function at zero time lag, Γ = *γ*^*noise*^(0).

### Measures of pair-wise interactions

Pair-wise measures are computed for pairs of neurons from the same recording session and are then pooled across sessions.

#### Correlation of trial-to-trial variability

We use *z*-scored spike counts, 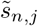. For each neuron, we define a vector of activation across trials, 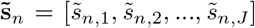. The correlation coefficient between activation of a pair of neurons *n* and *m* is computed as the Pearson correlation coefficient,

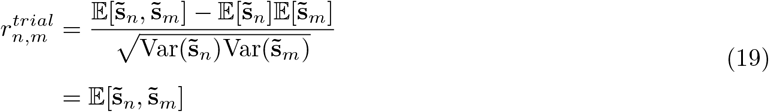

where the second equality is due to *z*-scoring [Bair et al., 2001].

#### Correlation of spike counts with binned spike trains

We bin spike trains in bins of length *L* = [20, 50, 75, 100] ms and count spikes in bins. Spike counts are *z*-scored for each bin independently,

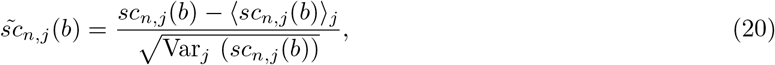

where *b* is the index of the bin, *b* = 1, 2, …, *B*, and the mean and variance are computed across trials. We define the vector of activation that concatenates binned spike trains across trials, 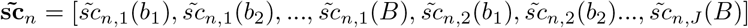. After this procedure is repeated for all neurons, we compute Pearson’s correlation coefficient between z-scored spike counts for each pair of neurons *n* and *m*,

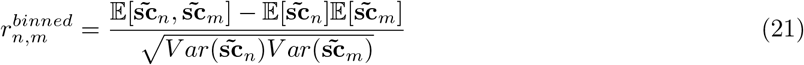

#### Synchrony

Now, let *n*, *m* ∈ {1, …, *N*} be fixed, and we define the spike trains of neurons *n* and *m* in trial *j* by *f*_*j*_(*t*_*k*_) := *o*_*n,j*_(*t*_*k*_) and *g*_*j*_(*t*_*k*_) := *o*_*m,j*_(*t*_*k*_). The cross-correlation function measures the co-occurence of spikes,

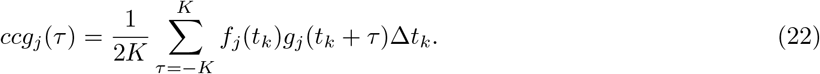

We also measure autocorrelation functions for neurons *n* and *m*. The autocorrelation function for neuron *n* is the following:

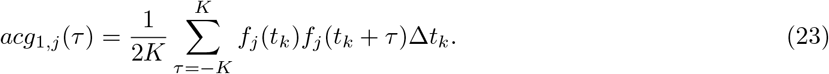

The cross-correlation function depends on the number of spikes of neurons *n* and *m*. To make it independent on the number of spikes, we normalize it with autocorrelations at zero time lag,

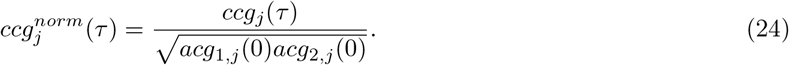

It is expected that an important part of the correlation is generated by the common input to the two neurons [Bair et al., 2001]. The correlation due to the common input is captured by the trial-invariant correlation function. The latter is computed from spike trains that are not from the same trial,

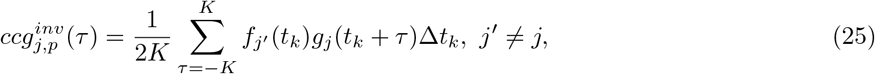

where *p* is the permutation index and *j*′ is defined with random permutation of the order of trials for the neuron *n*, while the neuron *m* keeps the correct order of trials. The permutation of the order of trials is iterated *nperm* = 100-times, and we average the resulting functions across permutations, to get 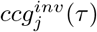. We capture the trial-specific correlation (i.e., the “noise” correlation) by subtracting the trial-invariant correlation from the overall correlation, and average across trials:

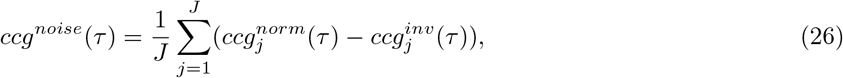

where *J* is the number of trials. The coefficient of synchrony is defined as the trial-specific cross-correlation function at zero time lag, *C* = *ccg*^*noise*^(0).

### Data and code availability

The dataset and the code generated during this study are freely available in a public GitHub repository: https://github.com/VeronikaKoren/struct_pop_cellreports. All results and figures reported in the article have been generated from the dataset and code in the repository. The code is published for documentation purposes. In case of questions, contact the lead contact Veronika Koren.

## Supplemental information

S1: Range of spike counts. Prediction accuracy during the target time window. Prediction accuracy during the test time window with a non-linear SVM.

S2: Decoding weights of the population model and the Area under the ROC curve of single neurons during the target time window.

S3: Weights of the population model during the first and the second half of the target time window.

S4: Synchrony within the group of informative and less informative neurons during the first and the second half of the test time window.

S5: Noise correlations of binned spike trains within the group of informative and less informative neurons during the target time window.

S6: Coupling and correlations within and across coding pools during the first and second half of the test time window.

