## supplemental information for "In V4, pair-wise synchrony and correlations depend on the structure of the population code"

### S1: Range of spike counts. Prediction accuracy during the target time window. Prediction accuracy during the test time window with a non-linear SVM.

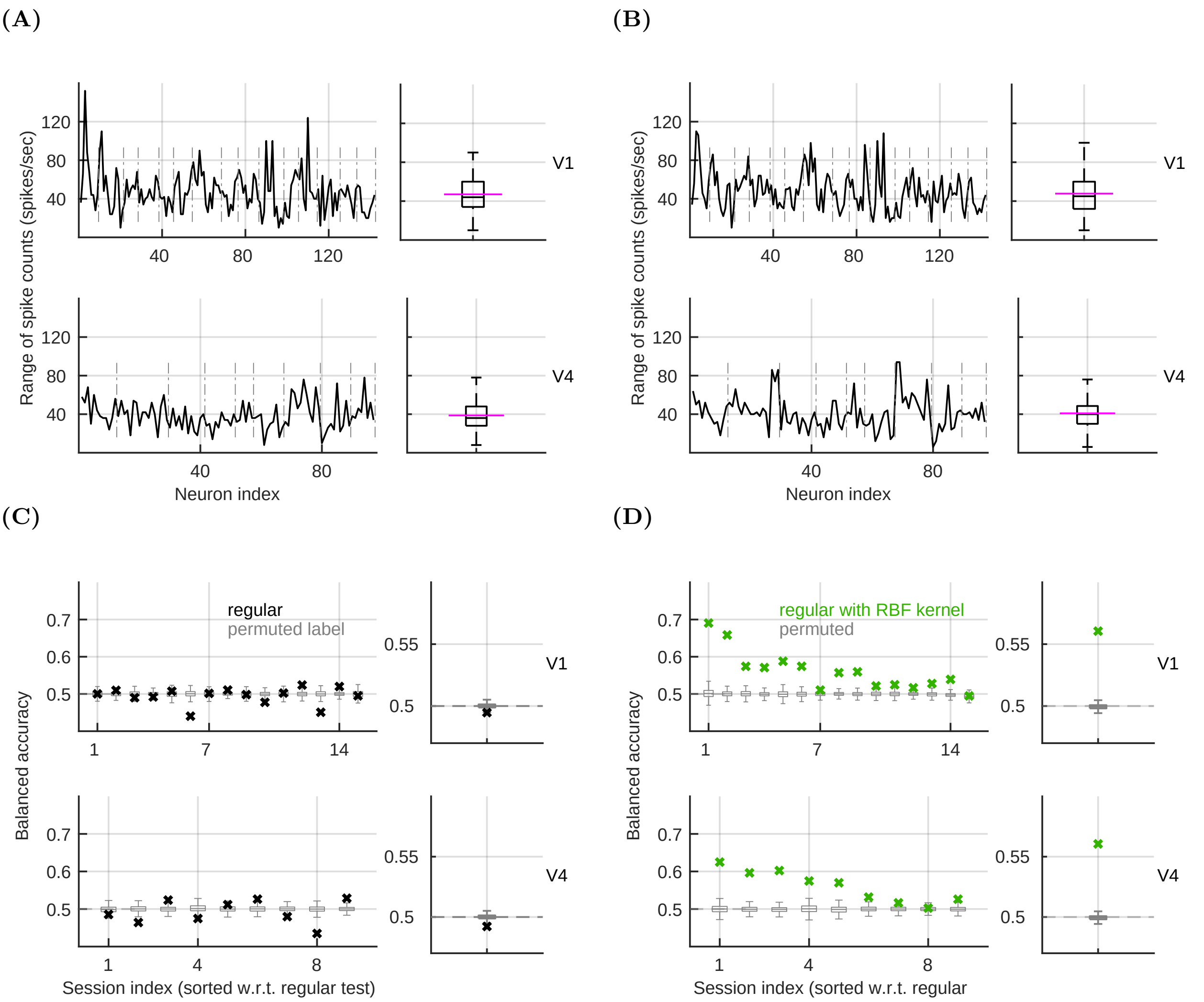


**Figure S1. During target, the prediction of the population model is at chance. A)** Range of z-scored spike counts during the test time window, in units of standard deviation. The range is computed for every single neuron, utilizing the distribution of spike counts across trials. We z-score the spike counts and compute the difference between the minimum and the maximum value. We use the entire time window ([0,500] ms w.r.t. the onset of the test stimulus). Left: The range of every single neuron, with dashed vertical lines delimiting sessions. The order of sessions is the same as in the Figure 1E. Right: Box-plot of the range. The magenta line shows the empirical mean. **B)** Same as in **A**, but for the target time window. **C)** Balanced accuracy of the population model (black) in V1 (top) and in V4 (bottom) during the target time window. Left: Prediction accuracy in recording sessions, sorted according to the performance of the regular model during test (Figure 1E). Gray boxplots show distribution of prediction accuracy of models with permuted class labels ($N_{perm} = 1000$). Right: Session-averaged prediction accuracy. The prediction accuracy during target is at chance. This is expected, since during target, the visual system does not yet have the information for discriminating matching and non-matching stimuli. **D)** Same as in **C**, but for the SVM with radial basis function kernel during the test time window. The prediction accuracy is similar to the linear SVM.

### S2: Decoding weights of the population model and the Area under the ROC curve of single neurons during the target time window.

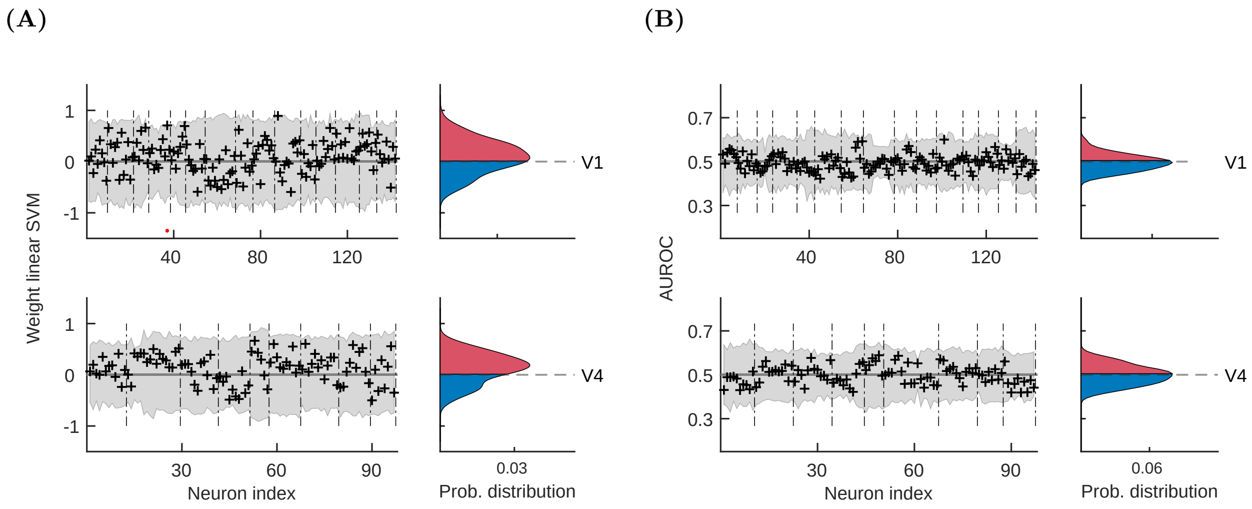


**Figure S2. During target, AUROC scores are centered around the chance value. A)** Left: Weights of the population model (black), collected across recording sessions in V1 (top) and in V4 (bottom). Dashed vertical lines delimit sessions. The gray area shows the distribution of weights of models with permuted class labels. Right: Distribution of weights, pooled across neurons from all recording session. **B)** Left: Area under the ROC curve (AUROC, black), collected across recording sessions. The gray area shows the distribution of results for models with permuted class labels. Right: Distribution of AUROC scores, pooled across neurons from all recording sessions. For both plots, we used $N_{perm} = 1000$ random permutations of class labels.

### S3: Weights of the population model during the first and the second half of the target time window.

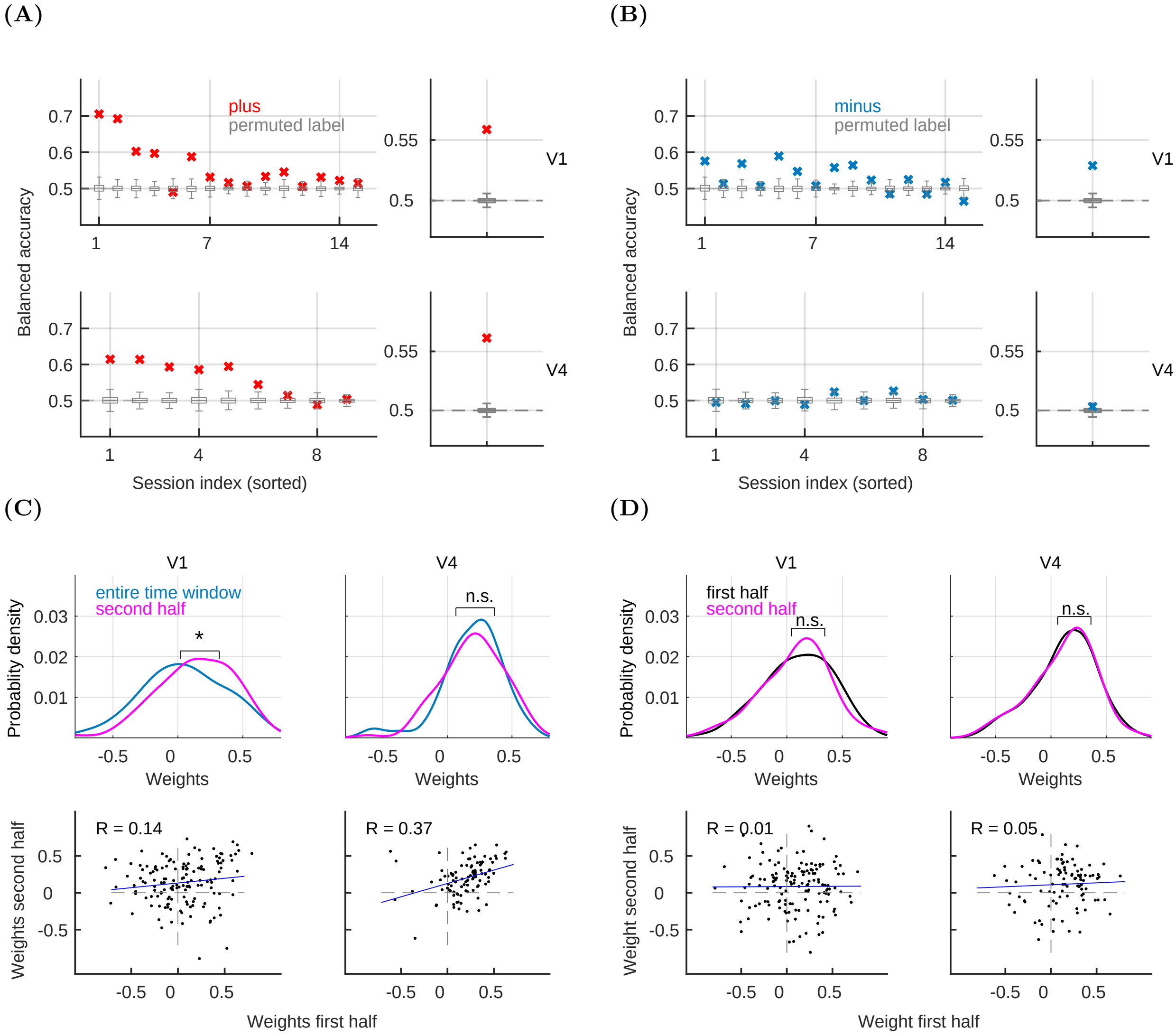


**Figure S3. Weights of the population model during the first and the second half of the target time window are uncorrelated A)** Prediction accuracy of neurons with positive weights in recording sessions (left) and averaged across sessions (right). The model uses all neurons from the recording session, but removes information from neurons with negative weights by replacing their activation with activation from randomly chosen trial. The trials of minus neurons are randomly permuted $N_{perm} = 1000$ times, without repetition. This way, the activity of minus neurons is associated with a random class label and minus neurons only contribute noise to the model. **B)** Same as in **A**, but showing the prediction accuracy of minus neurons, by removing the information from the activity of plus neurons. **C)** Top: Distribution of weights for the entire test time window (blue) and the second half of the test time window (magenta). We plot results in V1 (left) and in V4 (right). Asterisk marks significant change of the mean of the distribution (permutation test). In V1, the distribution during the second half is shifted towards positive weights with respect to the distribution of weights using the entire time window. Bottom: scatter plot of weights in the entire time window vs. the second half. $R$ is a linear correlation coefficient and the blue line is the least squares line. Weights using the whole and the second half of the time window are positively correlated. **D)** Same as in **C**, but comparing weights during the first and the second half of the target time window. There is no difference in distributions, and, interestingly, weights in the first and the second half of the time window are uncorrelated in both brain areas.

### S4: Synchrony within the group of informative and less informative neurons during the first and the second half of the test time window.

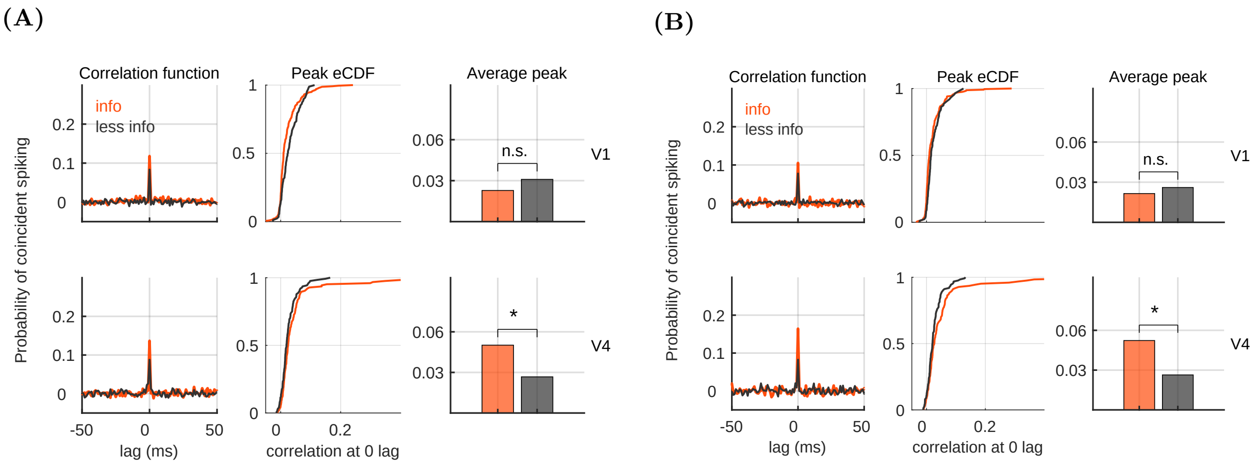


**Figure S4. Informative neurons in V4 are more strongly synchronized than less informative neurons. A)** Synchrony between pairs of informative and less informative neurons during the first half of the test time window ([0,250]ms w.r.t. the stimulus onset). Left: Correlation function for an example pair of informative (orange) and less informative (black) neurons. Middle: Empirical cumulative distribution function (CDF) of the correlation function at zero lag for informative and less informative pairs. Correlation function at zero lag measures synchrony of spikes. Right: Average synchrony for the two groups of neurons. Asterisk marks significance (permutation test with $N_{perm} = 1000$ random permutations). **B)** Same as in **A**, but for the second half of the test time window ([250,500] ms w.r.t. the stimulus onset). In both first and second half of the test time window, in V4, synchrony among informative neurons is stronger than among less informative neurons.

### S5: Noise correlations of binned spike trains within the group of informative and less informative neurons during the target time window.

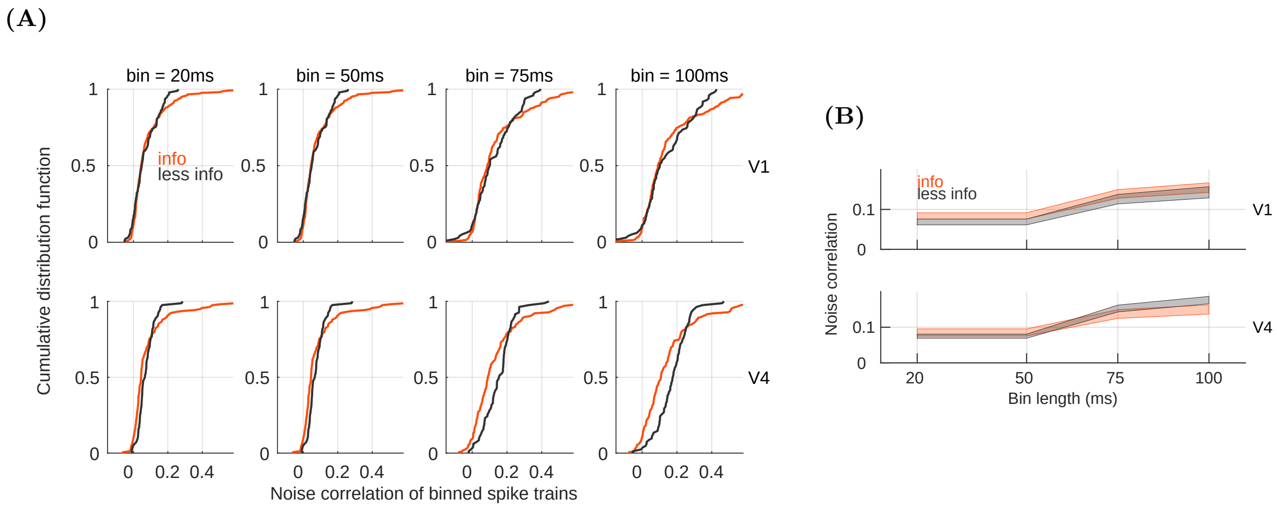


**Figure S5. During target, there is no difference in correlation strength between informative and less informative pairs. A)** Noise correlations of binned spike trains for informative and less informative pairs during the target time window. We show the empirical CDF for each group, measured in bins of [20,50,75,100] ms (plots from left to right). Correlations are measured between neurons from the same session, and are then collected across sessions. **B)** Mean ± SEM for results in **A**. Informative and less informative neurons do not differ in the correlation strength during the target time window (permutation test). All permutation tests use $N_{perm} = 1000$ random permutations.

### S6: Coupling and correlations within and across coding pools during the first and second half of the test time window.

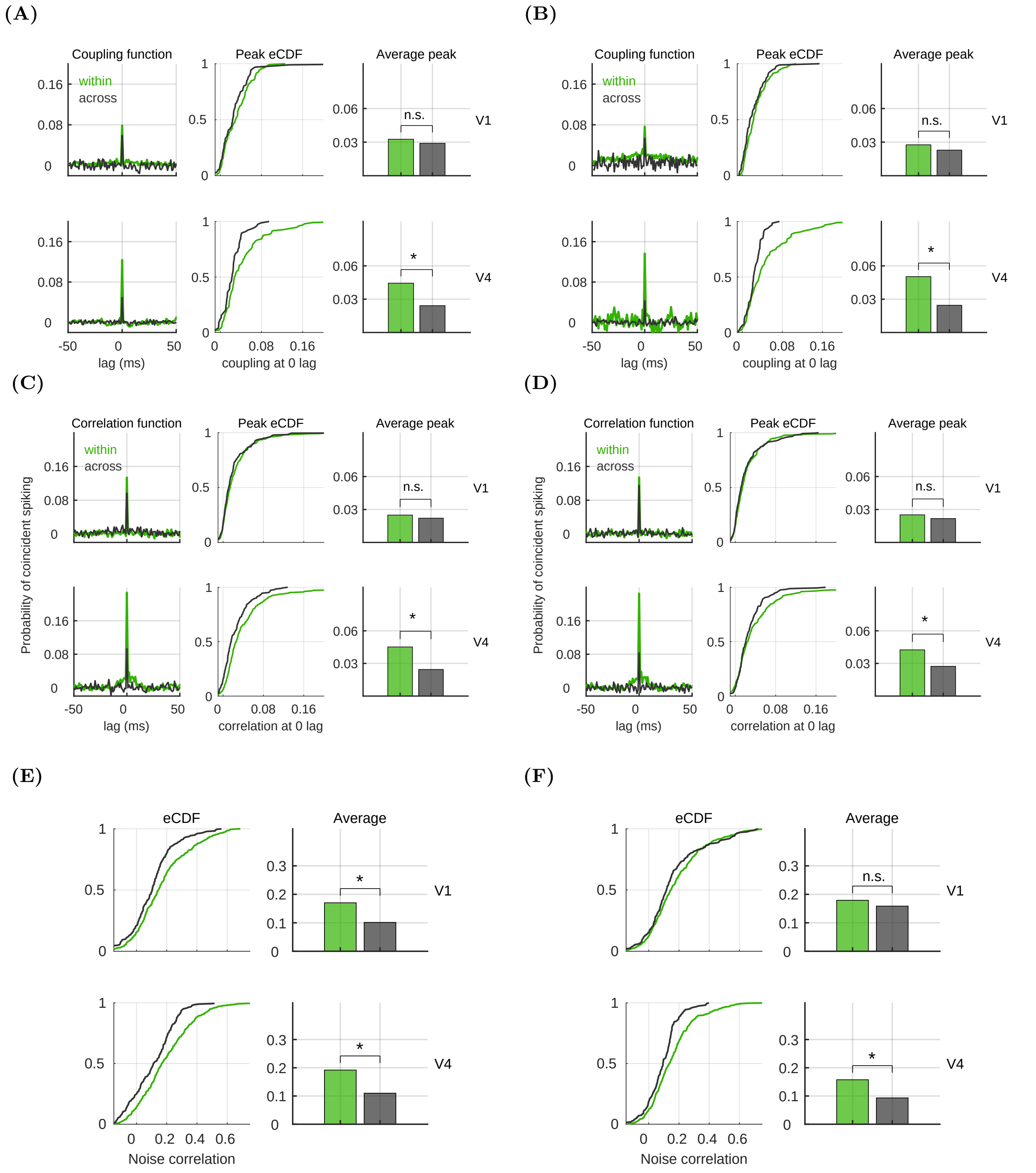


**Figure S6. Coupling, synchrony and correlations within pool are stronger than across pools. A)** Coupling within and across pools during the first half of the test time window ([0,250]ms w.r.t. the onset f the test stimulus). Left: The coupling function for an example neuron, for coupling within the pool (green) and across pools (black). The coupling function is plotted as a function of the time lag. Middle: Empirical CDF for coupling at 0 lag, distinguishing coupling within and across pools. Right: Average coupling at 0 lag, with asterisk marking significance (permutation test).

**Figure S6. B)** Same as in **A**, but for the second half of the test time window ([250,500] ms w.r.t. the onset of the test stimulus). **C)** Pair-wise synchrony within and across pools during the first half of the test time window. Left: Correlation function for two example pairs, one within and one across pools. Middle: Empirical CDF of the correlation function at zero lag, measuring synchrony of spiking. Right: Average synchrony. Asterisk marks significance (permutation test). **D)** Same as in **C**, but for the second half of the test time window. **E)** Noise correlations of trial-to-trial variability during the first half of the test time window. We show the empirical CDF (left) and the average correlation coefficient (right). Asterisk marks significance (permutation test). **F)** Same as in **E**, but for the second half of the test time window. For all plots, permutation tests use $N_{perm} = 1000$ permutations.
